# Programmable soft DNA hydrogels stimulate cellular endocytic pathways and proliferation

**DOI:** 10.1101/2024.05.26.595930

**Authors:** Ankur Singh, Nihal Singh, Manasi Esther Jinugu, Prachi Thareja, Dhiraj Bhatia

## Abstract

Hydrogels are pivotal in tissue engineering, regenerative medicine, and drug delivery applications. DNA molecules stand out among various biomaterials due to their unparalleled precision, programmability, and customization. In this study, we introduce a palate of novel cellular scaffolding platforms made of pure DNA-based hydrogel systems while improving the shortcomings of the existing platforms. DNA strands can form complex supramolecular branched structures essential for designing novel functional materials by its precise sequence-based self-assembly. These unique geometric scaffolds offer a soft, cushiony platform, ideal for culturing cells to mimic the complex native in vivo environments better. Each hydrogel comprises repeating units of branched DNA supramolecular structures, each possessing a distinct number of branching arms. The epithelial cells grown over these hydrogels show dynamic changes at multiple levels, from morphology to protein expression patterns, enhanced membrane traffic, and proliferation. The DNA hydrogels explored here are mechanically weak and soft and thus appropriate for applications in cell biology. This research lays the groundwork for developing a DNA hydrogel system with a higher dynamic range of stiffness, which will open exciting avenues for tissue engineering and beyond.

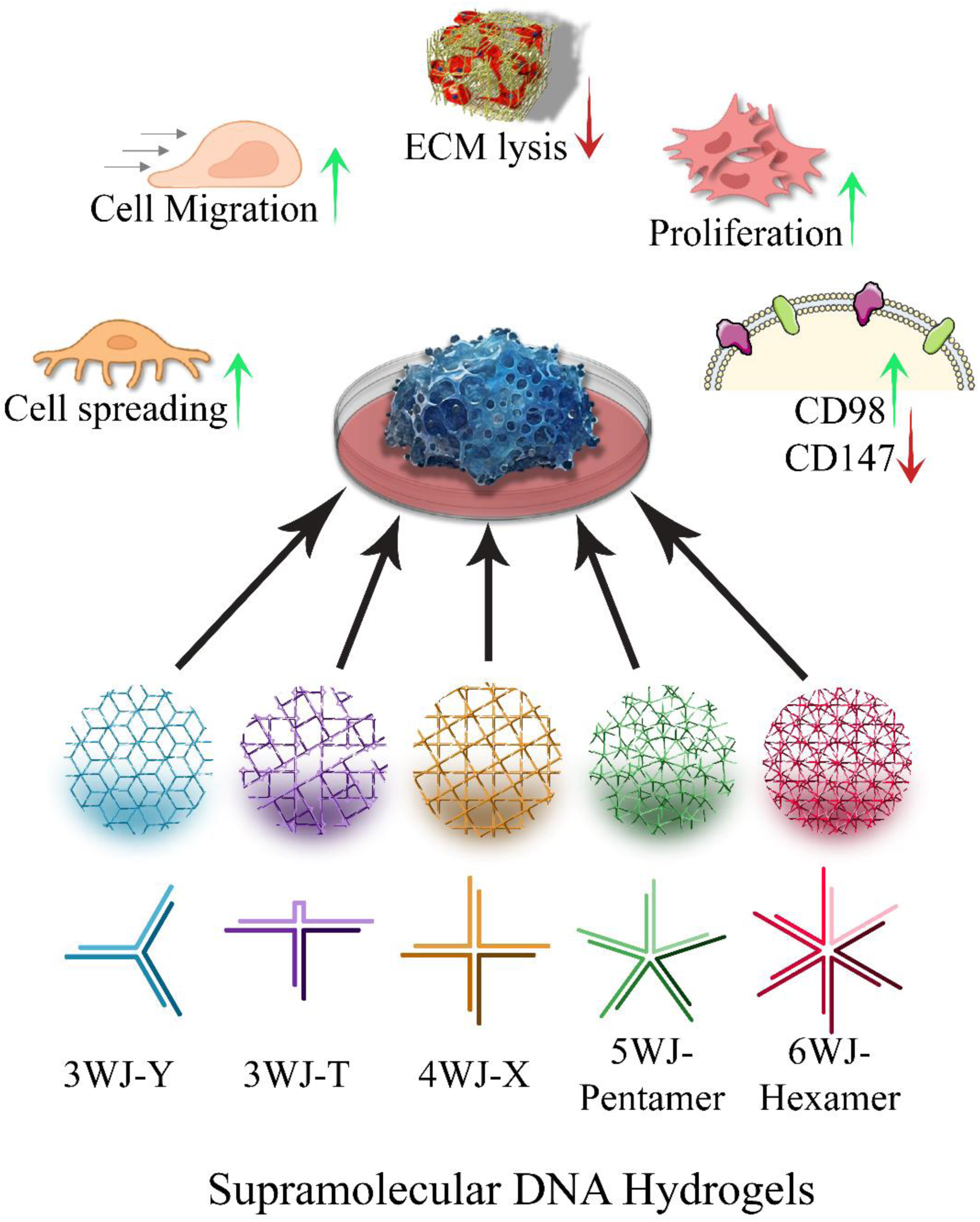

Graphical abstract illustrating diverse branched DNA supramolecular architectures forming DNA hydrogels of various geometric profiles, each put to use in the cell culture applications.

## Introduction

Recent times have seen a lot of significant milestone developments in the field of mechanobiology, especially in the context of understanding the key players in the mechanotransduction pathways. These mechanotransducers invoke mechanical stimuli-based perturbations within the cells, thereby modulating the cellular behavior^1–5^. These findings and developments have enabled us to appreciate the vital role of the Extracellular Matrix (ECM) and its beneficial properties toward the growth and maintenance of cells in their native environment. ECM is a heterogeneous mixture of fibrous proteins, glycoproteins, and proteoglycans. These components give rise to a scaffold in which cells attach, grow, mature, and differentiate. The crosstalk between ECM and the cells is highly dynamic and changes per the external stimulus. The ECM provides structural support and organization to the cells and plays an important role in vital cellular processes such as cell migration, proliferation, metabolism^6^, cellular respiration, cell-cell communication, and differentiation^7^. Depending upon the specific cellular need, the composition and distribution of the ECM are strictly regulated by the cells^1,8^. To date, the 2D monolayer culture has been extensively used for the majority of drug delivery applications, therapies, and disease model research. Nevertheless, the characteristics, appearance, functions, and biological processes of cells grown on a 2D flat surface are not precisely the same as those of an intended live animal system. The 2D cultured cells show flattened morphology, abnormal cellular polarization, altered response to pharmaceutical agents, and loss of differentiated phenotypes. These cells do not represent the original parental genotypic/ phenotypic variant, which strongly advocates the application of ECM-mimicking scaffold materials^9–11^.

To supplement structural and functional support for the cells grown in-vitro, various choices have been introduced, including growing the cells over patterned glass surfaces, elastomeric films, ceramics, foams, 2D fibrous coatings (poly-L-lysine), decellularized ECM scaffolds^12^, Hydrogels and others^13–15^. In the past two decades^16–24^, the application of hydrogels to deliver an ECM-mimicking environment for the cells in lab conditions seems to be the most optimal solution because of its high water-retention capability, tissue-like mechanics, and porosity^16,18,19,25–28^. Hydrogels are 3-dimensional networks of polymer with a high degree of water-adsorbing capabilities. The polymeric network acts as a scaffold, and the hydrophilic environment facilitates the diffusion of gases and the availability of nutrients, and media to the cells in the culture^28^. There are many suitable polymeric candidates for hydrogel synthesis, but the choice of deoxyribose nucleic acids (DNA) becomes obvious due to its inherent biocompatibility and precise programmability. The DNA strands are bi-polyanionic and polyelectrolytic and thus absorb growth factors, nutrients, media, and gases, making it an excellent candidate for hydrogel fabrications^29,30^. The precise base pairing by H-bond formation between the nucleosides of two different strands of DNA allows sequence-dependent hybridization, forming higher-order supramolecular architectures^31^. There are also naturally available and well-studied secondary structures of DNA that have important biological roles, both in structural and functional aspects. These include G-quadruplexes, i-motifs, hairpin loops, and cruciform structures. In cohorts with smartly designed aptamers^32,33^, these secondary structures offer additional functionality to the DNA-Hydrogels, including stimulus-responsiveness^34^ against changes in pH ^35–38^, temperature, presence or detection of a specific molecular target^13,35,39,40^, metal ions^13,14,41^, or generation of efficient and smart molecular probes^42–45^. The level of unmatched control, versatility, and programmability^46–49^ and its undeniable biocompatibility^50–52^ and stability make the DNA Hydrogels an ideal polymeric scaffolding platform for 3D tissue culture^53^.

Depending on how the DNA strands are used to make a hydrogel, we can broadly classify DNA-based Hydrogels into three broad categories. First, pure DNA Hydrogels are formed from physically interlinked ultralong linear DNA molecules^54,55^. The hydrogels formed from the smartly aligned short oligomeric strands of DNA come under the second category of pure dendritic DNA hydrogels^56,57^. Finally, the last category includes hydrogels made by combining the strengths of DNA with desirable functionality from other polymers, thereby forming hybrid DNA hydrogels^13,58–63^. The second category of short-stranded DNA hydrogels could be tailored to reproduce a network that can retain its features even at the nanometer scale^56,64^. This creates opportunities for us to explore a DNA hydrogel-based platform that could precisely control the geometric morphological patterns of the scaffold at the nanometer scale while retaining the stiffness and the polymeric lengths uniform. Despite significant progress, the field of biomaterials has yet to deliver a suitable polymeric system that can offer all these features.

This study explores the design, fabrication, and characterization of short-stranded dendritic DNA hydrogel systems for cell culture applications. More specifically, we demonstrate the efficacy of soft, self-assembled supramolecular DNA hydrogels in enhancing cellular spreading, remodeling lipid profile, and mitochondrial reorganization. We also show changes in the endocytic uptake routes, cell-surface protein expression patterns, and the perturbation in the focal adhesion sites after culturing the cells over DNA Hydrogels. This study aims to explore the potential of DNA hydrogel systems by offering the convenience of fabrication, geometric pattern precision, mechanical stiffness programmability, and scope for further functionalization. We hope to provide a range of soft DNA-based Hydrogel systems that offer unique mechanical and morphological features, vital for enhancing cellular spreading and proliferation. These soft DNA hydrogels’ network density and geometric profiles can easily be tweaked per the cell’s specific requirements. Through this study, we will try to address the following questions: Does the scaffold geometry play any role in cell spreading and proliferation? Will there be any observed beneficial effect of nanoscale morphological patterns despite the weak mechanical stiffness of the scaffold? What happens to the Focal adhesion sites and the state of ECM remodeling in such soft Hydrogels? These DNA based soft hydrogels will represent a new scaffold mimicking some features of extracellular matrix for modulating cellular physiology with biological and biomedical applications in stem cells and regenerative therapeutics.

## II. Results and Discussion

### 1. Synthesis and characterization of DNA hydrogels

We have successfully designed, synthesized, and characterized a range of soft, self-assembled DNA supramolecular hydrogels from the adaptations and modifications of the previously reported literature^65,66^(**Figure 1a, Supplementary Figure 1**). The sequences reported in supplementary Table 1 have been enhanced to possess additional GC-rich regions to improve thermal stability and strengthen the supramolecular assembled structures. These hydrogels are formed by distinct branched supramolecular monomeric building blocks resembling branched junctions stabilized by magnesium chloride salts. A supramolecule made with three strands is called a Three-way Junction (3WJ), which includes a structure resembling either a Y or a T-shape. T-type 3WJ fundamentally differs from the Y-type due to a hairpin loop structure inserted within the strand T2, making it sterically form a ‘T’ monomeric unit instead of a ‘Y’ shape. We have also fabricated various DNA supramolecular hydrogels comprising four, five, and six-branched structures, thereby forming 4WJ (X-type), 5WJ (Pentameric-form), and 6WJ (Hexameric form) type supramolecular monomeric units, respectively. The strands of these supramolecular DNA hydrogels (namely T, Y, and X forms) are designed in such a way as to leave six nucleotide-long adhesive ends after hybridization. However, the strands forming 5WJ and 6WJ form supramolecular HGs, with nineteen strands of sticky adhesive ends forming a network. The sequences of 3WJ and 4WJ are palindromic, and all the strands possess similar sticky adhesive ends for linking. This enables the system to rapidly bind to form hydrogel once we pool all the strands. In such a simple DNA HG system, it is difficult to have control over the gelation process. To regulate the gelation behaviors of these hydrogels, we modified the sticky adhesive ends of the 5WJ and 6WJ only to form gels when both the linker arms were available, thereby distinctly categorizing them into A and B monomeric forms (Supplementary Figure 1).

**Figure 1:**
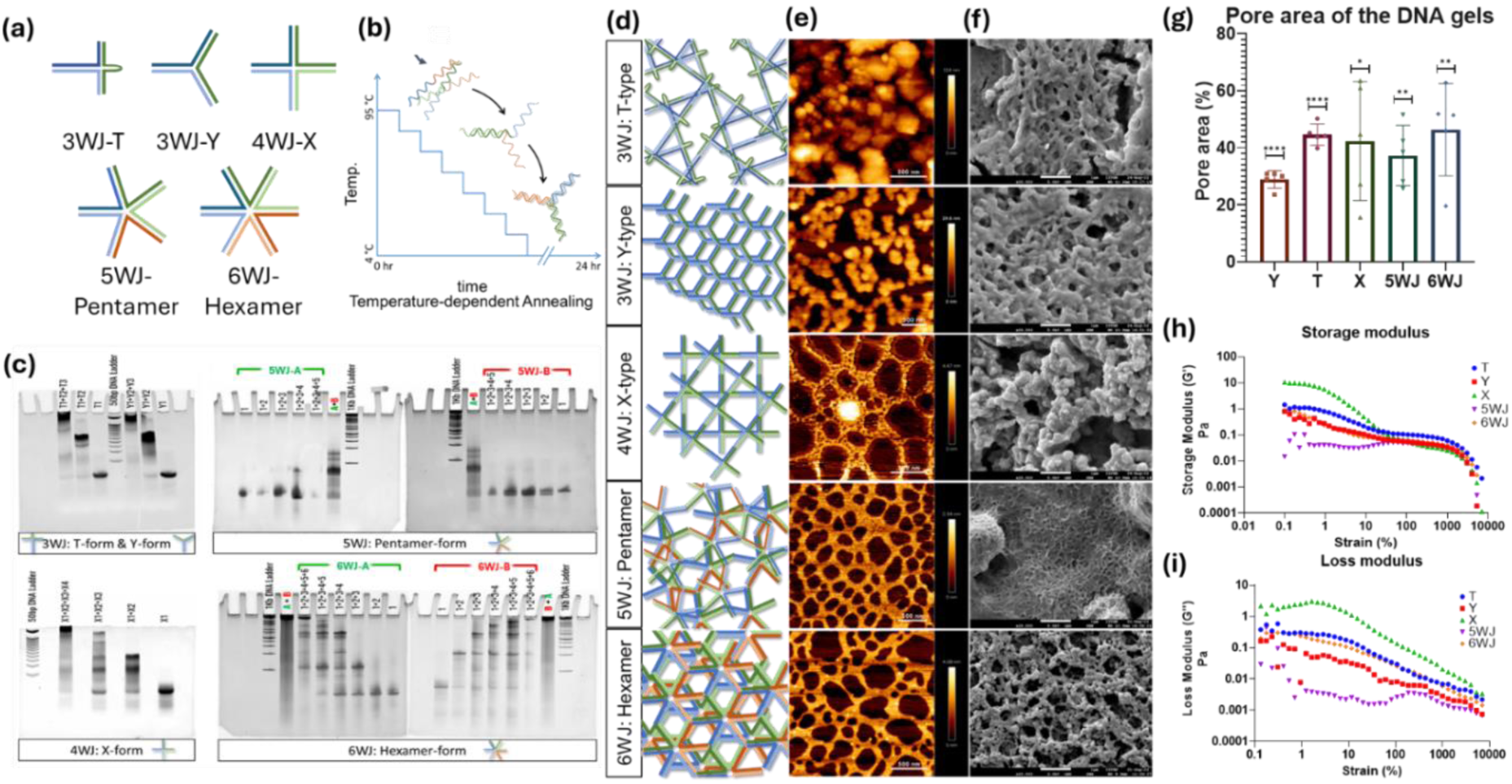
Synthesis and characterization of self-assembled DNA hydrogels. (a). schematics of various DNA supramolecular branched monomeric units. (b). Thermal annealing-based self-assembly of ssDNA is used to form higher-order branched monomers. (c). EMSA shows sequential binding of ssDNA and the shift in their electrophoretic mobility due to increased molecular weight. (d, e, f). shows graphical illustrations, AFM, and SEM images of DNA hydrogels at 50 µM concentrations, where scale bar = 500nm (e) and 1µm (f), respectively. (g). Pore area calculations based on thresholding of the void area (darker background) in the AFM and SEM images of the DNA hydrogels. ** signifies p-value of 0.0012, Welch’s ANOVA test. (h, i). DNA hydrogels ’ storage and loss modulus values were taken at 1Hz oscillating frequency when a strain sweep test was performed.

All five hydrogels were formed by sequence-based hybridization using a thermal annealing process (**Figure 1b, Supplementary Figure 1**). The formation of higher-order complexes was confirmed by observing the electrophoretic mobility shift in the band patterns of incrementally added strands of DNA. On a 10% non-denaturing native PAGE gel, the DNA hydrogel strands were loaded in equimolar ratio in an incremental order of strands added to the subsequent samples. For example, In a 3WJ T-type DNA Hydrogel, sample one comprises only T1 strands, while samples two and three will contain an equimolar ratio of T1 & T2, and T1, T2 & T3, respectively. (**Figure 1c, Supplementary Figure 2**). Because of the increase in molecular weight between sample three and sample two, we observe a shift in the EMSA band patterns. All DNA supramolecular Hydrogels showed ladder-like band patterns, indicating the hybridization of ssDNA to form a branched DNA building block necessary for forming DNA hydrogels. The formation of these higher-order complexes was also confirmed by running equimolar samples with varying ratios of the ssDNA forming each monomer in the Nanoparticle Tracking Analyzer (NTA) instrument (**Supplementary Figure 3)**. The samples containing the strands for 3WJ T-type Hydrogels in equimolar ratio of T1, T1+T2, and T1+T2+T3 strands showed a subsequent increase in the area, volume, and concentration of the higher-order complex being formed.

The structural morphology of the DNA hydrogels was performed by BioAFM and Field Emission Scanning Electron Microscope (FE-SEM). The AFM and SEM images showed an increased network density and well-defined morphological features of network structure from 3WJ to 6WJ DNA hydrogels (**Figure 1e,f; Supplementary Figure 4a, 5a**). Figure 2-d illustrates the graphical representation of possible 3D self-assembly of the supramolecular monomers to form complex network systems. The increment in morphological features and network density in association with the simpler to complex branched DNA supramolecular hydrogel systems were within our expectations. Our observations from simple to complex branched HG systems showed an increment in pore area of the hydrogel, a desirable quality in designer Hydrogel systems. The pore area of the dried hydrogels was calculated using the thresholding method applied over the darker regions (void spaces) on the AFM and SEM images (**Figure 1-g; Supplementary Figure 4b, 5b**). The Oscillatory Strain Sweep test was performed to test the visco-elastic behavior of these DNA hydrogels on a rheometer (**Figure 1h,i, Supplementary Figure 6**). At 1% strain and 1Hz oscillation, the ratio of G’ (Storage Modulus) to G” (Loss Modulus) for 3WJ-T, 3WJ-Y,4WJ-X, 5WJ-Pentamer, and 6WJ-Hexamer came out to be 5.22, 2.29, 2.06, 3.08, and 3.35 respectively (**Supplementary Figure 6h**). A higher G’/G” ratio indicates higher mechanical strength. The hairpin-loop structure, within the 3WJ-T form, sterically confers its mechanical stiffness, as indicated by their high G’/G” ratio. The results also depict an increment in the mechanical stiffness in these HG systems depending upon their network densities (indicated by G’/G”) from simple 3WJ to more complex 6WJ.

**Figure 2:**
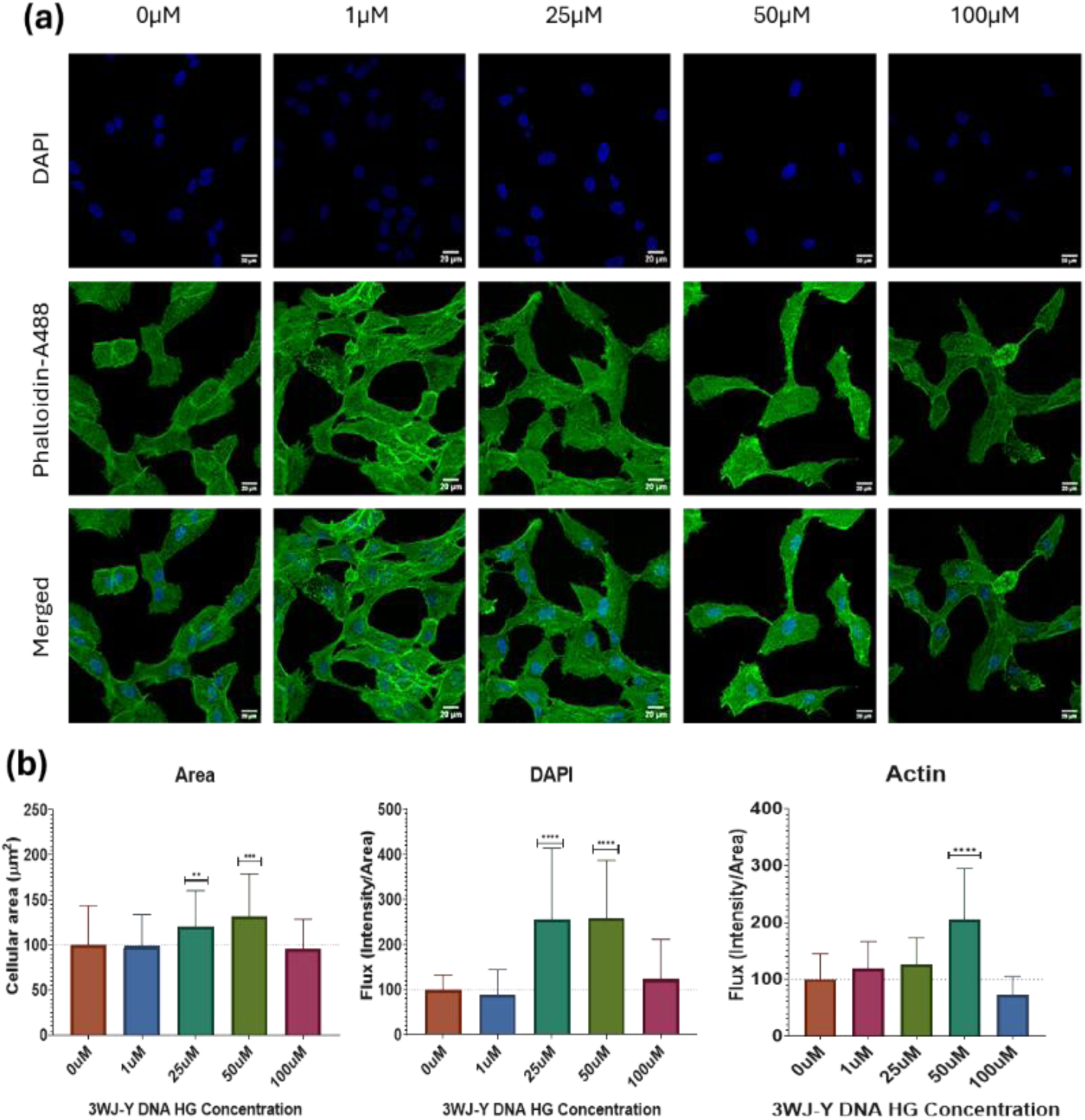
Effect of DNA Hydrogel concentration on cellular growth and spreading. (a) Confocal images of RPE1 cells grown over different Hydrogel Concentrations of 3WJ-Y type. The cells were grown over DNA hydrogel-coated coverslips for 24 Hours at 37°C; the Scale bar is 20µm. (b) Quantification of cells (n=30) shows cellular area, and flux (Intensity/ area) of DAPI and Actin, respectively, against various concentrations of 3WJ-Y type DNA hydrogels. ****, Statistically significant value of p<0.0001 (one-way ordinary ANOVA).

### 2. DNA Hydrogel stability

We performed the serum stability test for the stability of DNA Hydrogels in serum conditions (**Supplementary Figure 7**). We incubated all DNA hydrogels in 50% Fetal Bovine Serum (FBS) and tested the intensity of bands of DNA on 10% Native PAGE at an interval of 24 Hours for 3 days. The DNA bands with and without FBS showed no significant difference, indicating the stability of DNA hydrogels within 50% FBS solution for 3 days. All the cell culture studies conducted in this work were carried out in DMEM-complete media supplemented with 10% FBS and 1% antibiotics. Inferencing the serum stability results, the DNA hydrogels can remain stable for at least a week. The self-assembled supramolecular monomeric forms were stabilized by the presence of MgCl_2_ ions, raising the question of instability due to the presence of divalent cations within the cell culture grade DMEM media. To confer the effect of changing solvent in DNA HG synthesis, we tried culturing RPE1 cells over an HG-coated surface. If the DNA HG stability were compromised due to free divalent cations within DMEM-SF media, then the cells growing over such HGs would show some limiting growth features, such as diminished growth or proliferation. We prepared DNA HGs in both NFW (Nuclease Free Water) and DMEM-SF (Serum Free) media to test this. We cultured the cells over HG-coated coverslips, followed by fluorescently labeling the mitochondrial network and lipid bilayer to estimate cell mobility and spreading changes. (**Supplementary Figure 8a,b**). The effect of change in solvent for DNA HG preparation had no change when we switched from NFW to DMEM-SF media. To our surprise, there were no observable changes between both conditions, thereby confirming the stability of DNA hydrogels within the divalent ion-rich DMEM media.

### 3. Cellular adhesion and spreading

The adhesion of cells onto ECM-mimicking scaffolds is the first step in establishing an interaction between the ECM and the cells, further laying the foundation for cell regulation and cell-cell communication, which are crucial for cellular development and tissue maintenance^67–70^. **Supplementary Figure 9** shows a rough outline of the experimental plot, illustrating steps taken in prepping DNA HGs for culturing cells. The cells were then grown on DNA HG-coated coverslips and cultured in a CO_2_ incubator for 24 hours, then fixed and stained for confocal image acquisition. We tested the effect of change in polymeric concentrations in forming DNA HGs by culturing the RPE1 cells over 3WJ Y-type DNA-HG coated coverslips with varying concentrations ranging from zero (Control) to 100 µM of polymeric concentrations (**Figure 2**). Cell area calculations estimated the extent of cellular spreading. The cultured cells were stained with Alexa-488 fluorophore-tagged Phalloidin for actin labeling and DAPI (for the nucleus) as a counterstain. To our surprise, the highest concentration of 100 µM showed no significant difference in the cellular area compared to the control, which might be due to the abundance of negatively charged polymers of DNA within the Hydrogels (**Figure 2b**). However, smaller concentrations from 10µM to 50µM showed a substantial increase in the cellular area and the increase in the polymeric concentrations of DNA. The maximum cellular spreading capability was observed at the 50 µM Hydrogel concentrations; therefore, we will keep the hydrogel concentrations fixed at 50 µM in the subsequent cell-culture studies unless stated otherwise. We also noted a positive correlation between an increase in actin polymerization and an increase in the cellular area. Interestingly, we also observed a polymeric concentration-dependent increase in the intensity of the nucleus in the cells, indicating that the cells are rapidly dividing and have enhanced cellular spread. Our results suggest that by increasing the polymeric concentrations of DNA hydrogels, we can actively regulate the state of the cell during cellular adhesion and division. Similar observations were also noted in other DNA hydrogels.

### 4. Mitochondrial Expression and Lipid profile change

Observable changes within the cellular components, such as the composition and distribution of cell organelles, changes in the contents of lipids during cellular growth and division, and many more often accompany the rapid cellular division. To look into such changes resulting from DNA hydrogels, we grew the RPE1 cells over coverslips coated with various hydrogels. Coverslip without any coatings acted as our control. The RPE1 cells were stained with MitoTracker DeepRed and DIO dye to label mitochondria and lipid molecules within the cell membranes, respectively (**Figure 3**). Under normal physiological conditions, the mitochondria form a filamentous network, which spreads around the nucleus^71,72^. However, when the cells are cultured over soft scaffolding materials, it has been noted that the mitochondrial network undergoes fission. This is the cell’s response to adjust to the rapid cell division by focusing on segregating and sorting mitochondria instead of forming a filamentous network. Therefore, the mitochondria stay granular, as indicated by the decreased fluorescence of the MitoTracker DeepRed stain as compared to the control samples (Figure 4-a). Our results align with previously reported studies of culturing cells over soft collagen scaffolds^72^. We report a steady decrease in mitochondrial fluorescence signal in hydrogels of increased density, Highest in 3WJ T-type and lowest in the 6WJ Hexamer-type DNA hydrogels (**Figure 3b**, MT Raw Int., and Flux). Despite decreased MitoTracker expression, the cells remained viable and divided faster than the control group without showing signs of stress.

**Figure 3:**
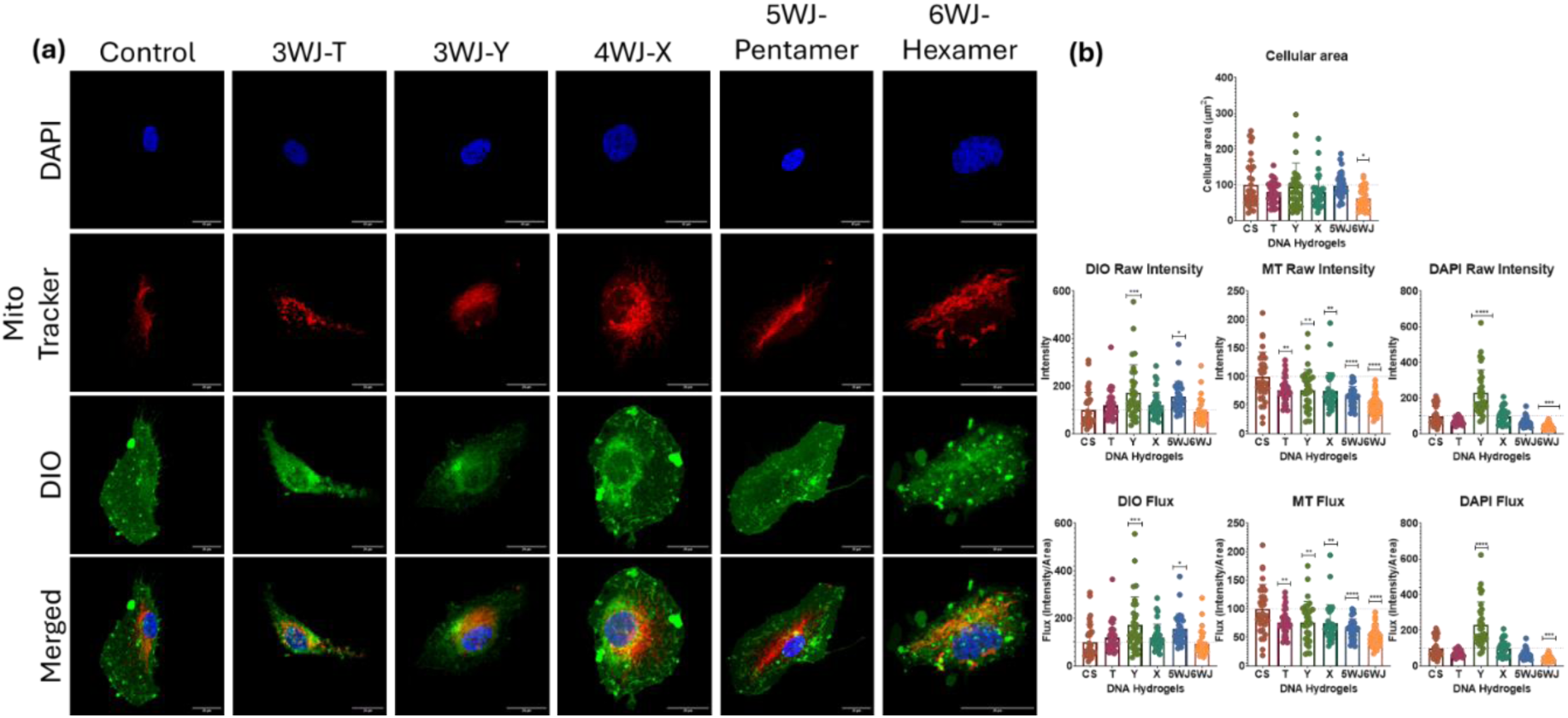
Effect of DNA Hydrogels on Mitochondrial assembly, distribution levels, and membrane lipid composition. (a) Confocal images of RPE-1 cells stained with DAPI (Blue channel), MitoTracker DeepRed (Red Channel), DIO (green channel), and merge channel, respectively. The DIO dye is highly lipophilic and stains the lipid bilayer membrane. Scale bar 20µm. (b) Quantified data of the cellular area (Top Row), Raw Intensity (Middle Row), and Flux (Intensity/Area) of DIO (Left), MT (middle), and DAPI (Right), respectively, with respect to changes in DNA Hydrogel systems. ****, statistically signifies p-values <0.0001; n=30 (One-way Ordinary ANOVA).

**Figure 4:**
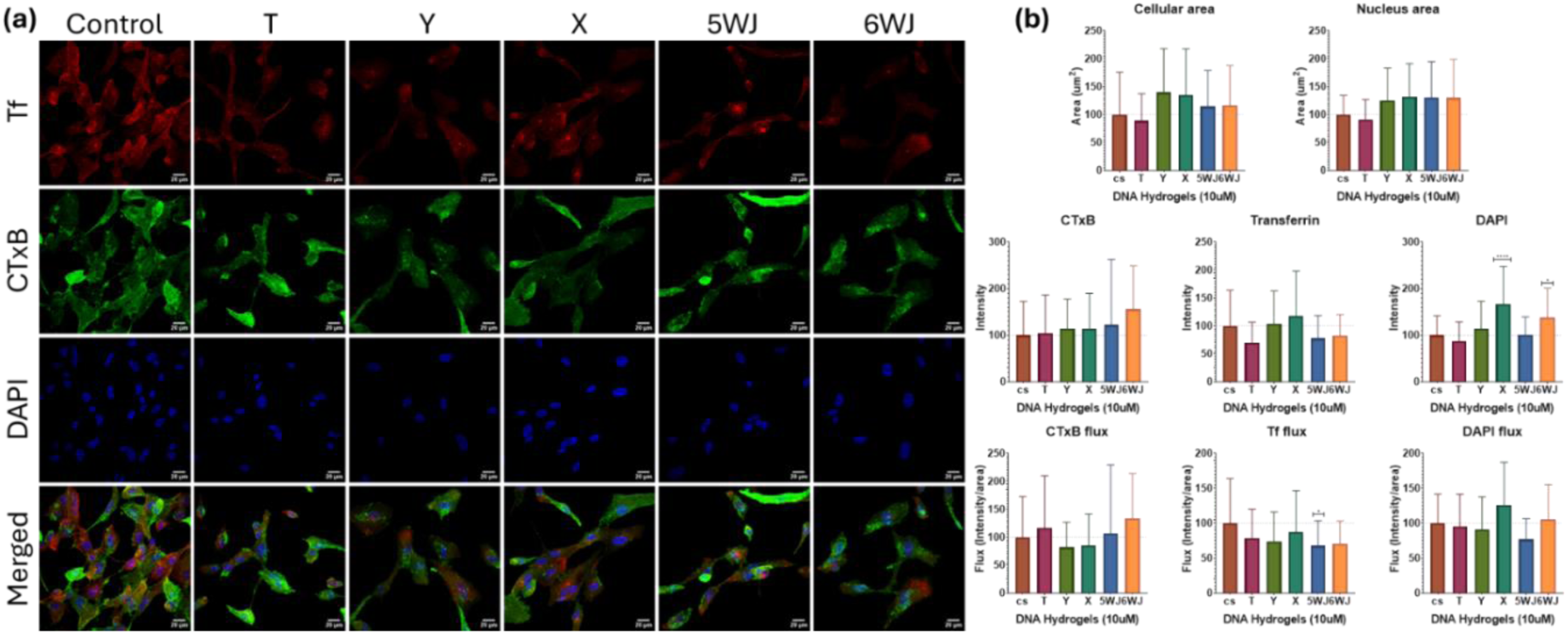
Endocytic Uptake study of Cholera Toxin-B (CTxB-FITC; Green), Transferrin (Tf-Alexa 647; Red), and DAPI (Blue) in RPE1 cells cultured over various DNA Hydrogels at 50 µM concentrations. (a) Confocal images showing the relative expression pattern of CTxB, Tf, and DAPI after 15 minutes of incubation post-treatment. Scale bar =20 µm. (b) Graphs showing quantified cellular area, nuclear area, Raw Intensity, and Flux of CTxB, TF, and DAPI, respectively. Statistics: **** signifies p-value <0.0001, n=30 (One-way ordinary ANOVA).

Conversely, the fluorescence signal of DIO-labelled lipid molecules has increased significantly in the order of increasing network density. Exceptionally, the 6WJ samples show no significant change in the fluorescence signal. This could be attributed to a fall in cellular area in the case of the 6WJ sample cells. High intensity of the DIO signal in other hydrogels shows an overall increase in the lipid content of the cells. To rule out the variation caused by the variable size ranges of the cells in consideration, we measured the flux (fluorescence intensity per unit area) of the fluorophore signals (Figure 4-b). By integrating the results of diminished MitoTracker DeepRed and increased DIO signals, we may conclude that the accelerated cellular division hinders the formation of mitochondrial networks and necessitates substantial input of new lipids to support the cellular division. However, more studies are needed to validate the relationship between scaffold-assisted cell culture and changes in lipid compositions.

### 5. Endocytic Uptake Study of Cholera Toxin B-submit and Transferrin

Enhanced cellular spreading and rapid cellular division are both highly energetic processes that can be met by switching to endocytic uptake processes. The cells cultured over the scaffold benefit from localized nutrient requirements and enhanced cellular spreading. The cultured cells therefore often tend to increase the nutrient uptake route or surface protein recycling via endocytic processes. The role of endocytic uptake processes in various cellular signaling transduction pathways has been well studied and documented in the past two decades^73,74^. Endocytic uptake processes are broadly categorized into Clathrin-mediated endocytic (CME) pathways and Clathrin-independent endocytic pathways (CIE). CME pathways are the most studied pathways responsible for the vesicular internalization of surface receptor proteins for their recycling or degradation^74^. To study whether there are any changes in the endocytic route, we selected two well-known proteins, namely Transferrin (Tf) and Cholera toxin B-subunit (CTxB), to map the endocytic pathway dynamics of RPE1 cells grown over various DNA hydrogels (**Figure 4a,b, Supplementary Figure 10a,b,c,d**). Our study shows an increment in the uptake of CTxB- a well-known marker for the CIE pathway, after 15 minutes post-treatment of CTxB to RPE1 cells cultured over DNA HGs of 50µM concentrations (**Figure 4a (Figure 4,b left: Top-CTxB Raw intensity, and bottom-CTxB flux)**). CTxB binds to well-studied GM1 glycosphingolipids on the cell surface and then gets internalized either via GPI-enriched endocytic compartments (GEECs) or Clathrin-independent Carriers (CLICs), where it follows retrograde trafficking to Golgi complexes or the endoplasmic reticulum^75–78^. Transferrin^79,80^, on the other hand, gets selectively uptaken in the case of 3WJ Y-type and 4WJ X-type hydrogel conditions (**Figure 4b middle: Top-Transferrin Raw intensity, and bottom-Transferrin flux)**). The endocytic uptake process usually takes 45 mins to recycle the uptaken cargo by fusing it with the lysosomes. We demonstrate the dynamics of the endocytosis uptake process of both CIE and CME at 15 minutes (Figure 5-a,b), 30 minutes (**Supplementary Figure 10a,b**), and 45 minutes (**Supplementary Figure 10c,d**) of post-treatment of CTxB and Tf, respectively. We find that RPE1 cells, when cultured over self-assembled DNA-based supramolecular Hydrogels, tend to prefer the CIE pathway rather than the CME pathway due to lower energy consumption in the case of CIE.

**Figure 5:**
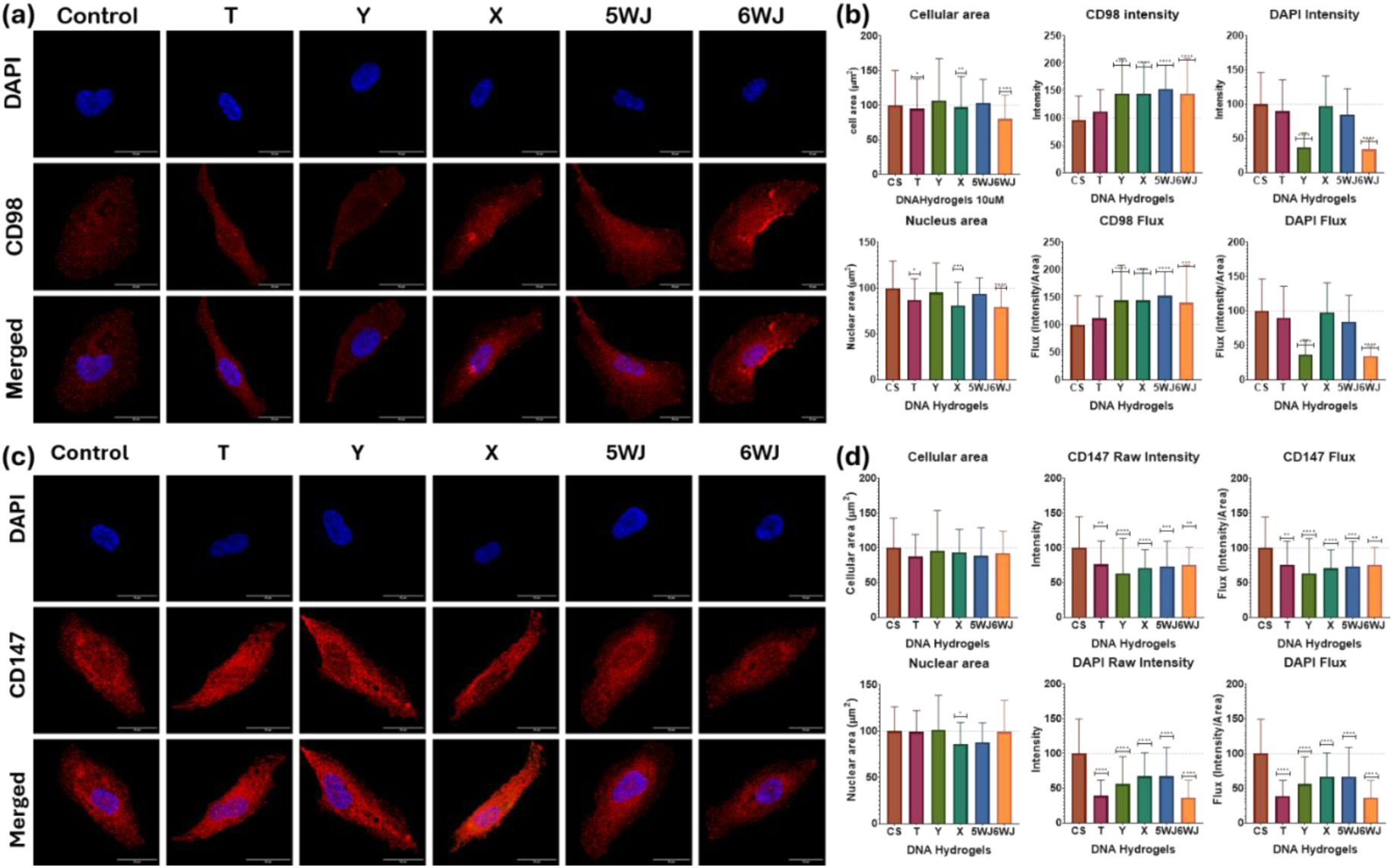
Immunofluorescence study of CD98 (a,b) and CD147 (c,d) expression levels in RPE 1 cells grown over 50 uM of DNA Hydrogels. (a) Confocal microscopic images of RPE 1 cells stained with CD98 (Red) and DAPI (Blue). The scale bar is 20um. (b) Quantified data showing changes in cellular area, nucleus area, Intensity, and flux changes of CD98 and DAPI, respectively. N=30 cells; **** signifies p-value< 0.0001 (Ordinary One-way ANOVA). (c) Confocal microscopic images of RPE 1 cells stained with CD147 (Red) and DAPI (Blue). The scale bar is 20um. (d) Quantified data showing changes in cellular area, nucleus area, Intensity, and flux changes of CD147 and DAPI, respectively. N=30 cells; **** signifies p-value< 0.0001 (Ordinary One-way ANOVA).

### 6. DNA Hydrogels Influence Membrane Receptor Expression

Apart from the general shift in the preference of less-energy-consuming endocytic route, the RPE1 cells also show perturbation in the expression pattern of cell-surface proteins such as CD98 and CD147. The CD98 is a transmembrane glycoprotein that has been involved in the regulation of integrin signaling, which is linked to the activation of the Erk-Ras-MAPK pathway, which regulates essential biological processes such as cell proliferation, differentiation, survival, migration, and others^81–83^. The CD147, on the other hand, is surface-bound matrix metalloprotease associated with interaction ECM lysis upon stimulation^84–87^. Activation of CD147 is often accompanied by induction of Matrix Metalloproteinases (MMPs) that regulate the degradation of ECM. The RPE1 cells grown over DNA hydrogels showed an increased propensity for energy-efficient processes to match up their demand for rapid cell division. We observed that the RPE1 cells grown over DNA HG-coated coverslips showed increased CD98 expression (**Figure 5a,b**) and decreased CD147 activity (**Figure 5c,d**) by the increasing network density of DNA HGs. Our results suggest that higher junction DNA HGs promote cellular growth, rapid cell divisions, proliferation, and migration of the cells. We also note that ECM degradation is halted by regulating MMPs, providing optimal conditions for the cells to grow and populate the DNA HGs.

Till now, we have demonstrated the surface receptor expression pattern of CD98 and CD147 at 50 µM DNA HG concentrations. However, we were curious to see the perturbation in the expression pattern at lower polymeric HG concentrations. We thus prepared lower concentrations of DNA Hydrogels, i.e. at 10 µM, the expression pattern of CD98 (**Supplementary Figure 11a,b**) and CD147 (**Supplementary Figure 11c,d**) falls drastically. The inference we obtained was inconclusive because of the lack of statistical power to conclude any positive or negative correlation. (**Supplementary Figure 11**).

### 7. DNA hydrogel stiffness regulates the Focal adhesion Complex

Culturing the cells over a scaffold-coated coverslip will trigger the cells to attain normal spherical morphology compared to flattened morphology in 2D monolayer cultures. The spherical cells interact with other cells or the matrix by localized points of protein complexes called Focal adhesion (FA) sites. To address the question of whether the stiffness or the geometry of the DNA hydrogels has any effect on the Focal Adhesion (FA) complexes, we monitored the expression of Vinculin, β-integrin, and Cadherin. Vinculin^88–90^ and β-integrins^81^ are the two most important and well-studied key proteins of the FA complex, thus regulating the cell-matrix interactions and are crucial for conducting mechanotransduction signaling across the Cell-ECM axis. E-cadherin, however, is involved in the mechanical coupling between two neighboring cells^91,92^. We observed striking decreased intensity and flux levels of Vinculin (**Figure 6a,b**) and β-integrin (**Supplementary Figure 12**), indicating the FA complexes are decreased in size and function to promote cellular mobility in the 3D DNA Hydrogels in the order of increased network density. The DNA HGs presented in this work are all mechanically weak compared to other pure DNA HGs. The E-cadherins are all upregulated in the case of DNA HG except the 3WJ Y-type. The highest expression of E-cadherin was observed in the 6WJ DNA HGs (**Supplementary Figure 13**). These results indicate a basal level of cell-cell adhesion phenomenon taking place while the cells actively divide, migrate, and occupy the 3D milieu within these DNA HGs. Notably, the trend follows the increasing order of DNA HG network densities. However, further experiments are needed to accurately validate the number and the size of FA sites diminished upon cellular growth over soft DNA HG samples.

**Figure 6:**
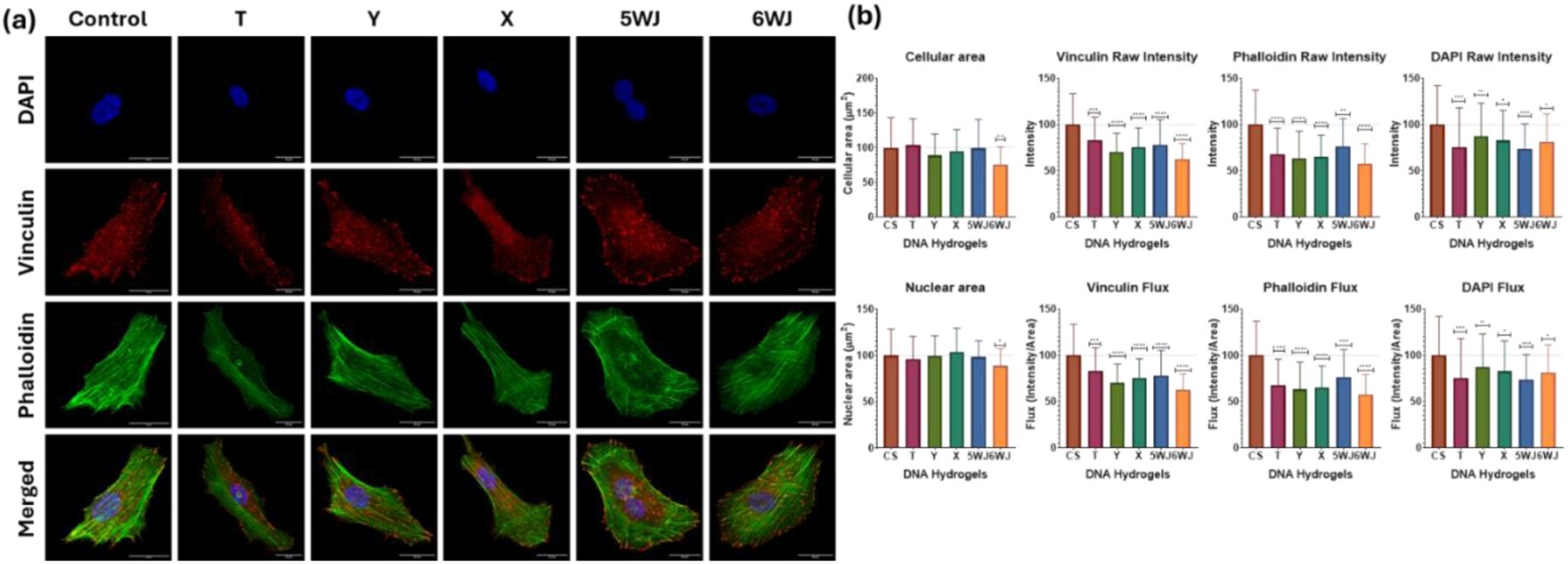
Immunofluorescence staining of Vinculin, phalloidin, and DAPI in RPE1 cells grown over 50uM of DNA Hydrogels. (a) Confocal microscopic images of RPE 1 cells stained with Vinculin (Red), Phalloidin (Green), and DAPI (Blue). The scale bar is 20um. (b) Quantified data shows changes in the cellular area, nucleus area, intensity, and flux of Vinculin, Phalloidin, and DAPI, respectively. N = 30 cells; ****, signifies p-value < 0.0001 (Ordinary One-way ANOVA).

## III. Conclusions

We synthesized self-assembled DNA Supramolecular Hydrogels of increased network densities while keeping their mechanical stiffness low. These types of soft hydrogels have their undisputed utility in many biological applications, such as drug delivery platforms for tumors, and show excellent biocompatibility, biodegradability, and programmability. The impact of varying Hydrogel network density on cell behavior in an in-vitro cell culture system has not been extensively studied before. We have selected DNA-based supramolecular assembly to answer this question to generate hydrogel systems of similar inter-nodal length but different network densities, thereby tweaking the geometry of the HG network system, thus formed. The Hydrogel with the smallest network density was 3WJ Y-type, followed by 3WJ-T type, 4WJ, 5WJ, and 6WJ being the densest. EMSA confirmed the formation of these sequence-dependent supramolecular assemblies. Morphological characterization of these dried hydrogels reveals a density-dependent structural assembly, as confirmed by AFM and SEM images. These images further helped decipher the total pore area of these hydrogels, which is the smallest in the case of 3WJ (Y) and largest in the case of 6WJ. A strain sweep test at 1Hz oscillating frequency in a rheometer revealed the poor mechanical performance of these hydrogel systems. However, the G’/G” ratios roughly follow a similar trend of having less mechanical stiffness in 3WJ (Y) and higher stiffness as we go from lower to higher-density Hydrogels. the 3WJ (T) hydrogels showed an exceptionally high G’/G” ratio, which can be explained by its sterically confined arms in 3D spatial orientation.

Before testing these self-assembled DNA HGs, we checked the stability of these HG systems under harsh serum conditions (50% v/v) and divalent cation-rich solvents used in cell cultures. We observed phenomenal stability for 72 hours at 50% serum conditions and no significant changes with the change in solvents.

These supramolecular DNA Hydrogels showed a concentration-dependent cellular spreading behavior, showing a 50% increase in cellular area and a 100% increase in actin-polymerization states at 50 µM of DNA HG concentrations (3WJ Y-type). We demonstrated that the DNA HGs at 50µM concentrations showed the best cellular spreading and actin-polymerization capabilities. The rest of the cell-culture experiments were conducted at 50 µM concentrations. Cellular studies showed that the DNA Hydrogels, in the order of their increased network densities, increased cellular spreading, mitochondrial fragmentation, lipid content, CIE pathway, CD98 intensity, and E-cadherin expression profiles. The pathways that showed downregulation included the CME, CD147 activity, Vinculin, and β-integrin levels. The overall cumulative effect of denser DNA HGs includes enhanced cellular spreading, division, proliferation, and clathrin-independent endocytic uptake of cargoes.

These soft scaffolding Hydrogels provide a repertoire of cell-culturing platforms intended to program cellular behavior by its conserved spatial geometric assembly patterns, which can be extensively used as a drug delivery platform against tumor model systems. We have successfully demonstrated the ease of synthesis of these hydrogels, varying in shape, size, and density while retaining excellent biocompatibility, biostability, precise programmability, and scope for further functionalization. The self-assembled DNA supramolecular Hydrogel is highly versatile and well-suited for potential use in real-world applications, including but not limited to 3D cell culture, tissue engineering, therapeutic purposes, studying complex disease models, and serving as a valuable platform for understanding the complex processes of mechanobiology.

## IV. Materials and Methods

All oligonucleotides (Table 1) were purchased from Merck (Sigma-Aldrich), at a 0.2µM synthesis scale with desalting purification. All oligonucleotides (Table S1) at a 0.2 μM synthesis scale with desalting purification were purchased from Merck (Sigma-Aldrich). Moviol, Triton-X, loading dye, Alexa Fluor (phalloidin) 488, moviol, and Hoechst (DAPI) were also purchased from Merck (Sigma-Aldrich). Ammonium persulfate, ethidium bromide, Triton-X, tetramethylethylenediamine (TEMED), paraformaldehyde, and cell culture dishes for adherent cells (treated surface) were procured from Himedia. Dulbecco’s modified Eagle’s medium (DMEM), fetal bovine serum (FBS), penicillin−streptomycin, and trypsin−ethylenediaminetetraacetic acid (EDTA) (0.25%) were purchased from Gibco. Tris-acetate-EDTA (TAE), and acrylamide/bisacrylamide sol 30% (29:1 ratio for SDS PAGE) were procured from GeNei. Magnesium chloride, NaCl, KCl, Na_2_HPO_4_, and KH_2_PO_4_ were purchased from SRL, India, and Santa Cruz Biotech. The Cholera Toxin B subunit-FITC Conjugate, Transferrin-Alexa Fluor 647, were purchased from Merck (Sigma Aldrich). E-cadherin rabbit mAb, Integrin β-1 Ab, and Anti-mouse IgG (H+L) F(ab)2 Fragment (A647) were purchased from Cell Signaling Technology. MitoTracker Deep Red FM, Phospho-vinculin (Tyr100) polyclonal Ab, Goat anti-rabbit iGg(H+L) cross-adsorbed secondary antibody Alexa-568, Phalloidin Alexa-647, and Vybrant Di0 cell labeling solution was purchased from Invitrogen (Thermo Scientific). CD44-FITC mouse mAb was purchased from Abcam. CD147 mouse mAb and CD98 mouse mAb primary antibodies were purchased from Biolegend.

1. **Cell culture.** Human retinal pigment epithelial-1 cell lines (RPE-1 cells) were obtained as a gift from Prof. Ludger Johannes at Curie Institut, Paris, France. RPE-1 cells were cultured in Dulbecco’s modified Eagle medium (DMEM) supplemented with 10% Fetal Bovine Serum (FBS) and 100 IU/mL penicillin−streptomycin (Penstrap) at 37⁰C with 5% CO_2_ and 95% Relative Humidity (RH) levels. The studies were performed in Phosphate Buffer Saline (PBS) of strength 1x and pH7.4 and serum-free DMEM media.

2. **Preparation of DNA supramolecular networks.**

All the monomeric oligonucleotides were made to self-assemble in a temperature-controlled annealing process to form a DNA supramolecular network of defined Junctions. A junction made up of three, four, five, and six single-stranded Oligonucleotides can form Three-way (‘Y’ and ‘T’ forms), Four-way (‘X’ or ‘Plus’ form), Five-way (‘Pentamer’), and Six-way (‘Hexamer’) junctions respectively. The 3’-ends of these oligonucleotides possess an extra stretch of sequences (sticky ends), which facilitates the binding with other junctions. These sequences have been adopted from previous studies^1,2^ and modified to improve the thermal stability of the sequences and enhance network formation capabilities via sticky end joinings (as shown in **Figure 1-a**). To efficiently control the network formation in higher-order junctions, we devised a strategy to join A-motif with B-motifs to form a fully connected network. Both the motifs share a common core region and vary in motif-specific sticky region, such that only A motif and B-motifs alone cannot form any network, while A+ B can undoubtedly form.

All oligonucleotides were mixed in an equimolar ratio of 100µM, which were stabilized by 2µM MgCl_2_ (as a bridging agent). The resultant mixture was heated at 95⁰C in a PCR machine and then gradually cooled down in decrements of -5⁰C/cycle, each cycle lasting for 15 mins. At the end of these reactions, the final product, Y, T, X, Pentamer, and Hexamer, is kept at 4⁰C for further studies.

3. **Characterization of DNA supramolecular networks.**

The DNA supramolecular network formation status, as well as its morphological features, was characterized by performing the following techniques.

### 3.1. EMSA

The classic tell-tale sign of confirming a strand-dependent structure formation is by performing the Electrophoretic Mobility Shift Assay. A 10% Native PAGE (Polyacrylamide Gel-Electrophoresis) was prepared in 1X TAE (Tris-Acetate-EDTA) buffer. Freshly prepared 2µL DNA samples were mixed with 1µL of 6X DNA-loading dye (having 0.25% Bromophenol Blue, 0.25% Xylene cyanol, and 30% Glycerol (v/v)). The mixed samples are then loaded within the lanes of the freshly prepared Native PAGE gel and then run at 90V for 90 minutes in 1X TAE buffer. The DNA bands were stained by Ethidium Bromide and were visualized in a Bio-Rad gel-documentation system.

### 3.2. Pore-area calculations

The AFM and SEM images were loaded onto Fiji (ImageJ 1.54f), and then a 30% thresholding algorithm was applied to highlight the void volumes in the DNA HG images. The highlighted area was used to generate a mask, with which we subtracted the original image file (8-bit format) to generate a composite image with only the void volume available for measurements. The Void volume, as well as the original images, were used to calculate the area occupied by the pores as well as the overall area of the HG, a difference of which returned the total pore area within these HG systems of different network densities.

### 3.3. NTA

The sequence-dependent network formation of DNA supramolecular architectures was confirmed by monitoring the change in the hydrodynamic radius of the higher-order complexes using a particle size analyzer instrument of the particle matrix. The instrument was calibrated using Polystyrene spherical beads (PS 100nm ∼5µg/L), and the laser to measure the particle size was 488nm laser. Samples of 1mL were prepared in a final concentration of 1µM and stabilized by 20nM of MgCl_2_. For a 3WJ-HGT type DNA scaffold, three tubes of varying oligomer count were prepared, such that the first tube contained only T1, the second tube contained T1+T2, and the third tube contained T1+T2+T3.

### 3.4. AFM

The samples were freshly prepared, drop-casted over a cleaved mica sheet, and dried overnight within a vacuum desiccator. The morphology of the dried DNA network was imaged by the Bruker NanoSense+ instrument, which was operated in tapping mode. The cantilever used was a gold-coated triangular RFESPG-75 cantilever with an 8-12nm tip radius and 75KHz frequency. The samples were imaged in 512*512 image format at various sizes, capturing the sample’s height, thickness, morphology, and porosity.

### 3.5. SEM

The samples were freshly prepared and drop-casted 10µL of DNA hydrogel samples over a clean carbon-coated stub followed by two-day lyophilization to remove water. The dried samples were sputter-coated with platinum atoms for 90 seconds before loading the sample into the vacuum chamber of the SEM instrument. The JEOL JSM-7900F field emission scanning electron microscope (FE-SEM) imaged the sample at an acceleration voltage of 5kV. The images were taken at several magnifications across all the samples.

### 3.6. Rheology

An oscillatory strain sweep test at 1Hz oscillation was performed on freshly prepared DNA HG samples via the Antonn Parr rheometer of model MCR302. A sand-blasted rough surface parallel plate geometry of 25mm was used to characterize the hydrogel samples. The experiment was performed at 41 distinct measurement points, ranging from 0.1 % strain to 7000%, at a constant temperature of 25°C. The hydrogel samples were taken at a 50 µM concentration for this experiment.

**4. In-vitro cell-culture studies**

The DNA samples were coated onto the 10mm coverslips by allowing them to adhere to the surface overnight within the wells of a 24-well plate. Five different DNA networks of 3WJ-Y, 3WJ-T, 4WJ-X, 5WJ-Pentamer, and 6WJ-Hexamer were taken for the comparative study. The pilot study on 3WJ-Y type was carried out in eight different concentrations, including 100 µM, 50 µM, 25 µM, 10 µM, 5 µM, 1 µM, 0.1 µM, and 0 µM (acting as our control). DAPI was used for staining the nucleus, Phalloidin/ Alexa488 was used for Actin labeling, DIO dye was used to label the phospholipids on the cell’s membrane, and MitoTracker DeepRed was used to label the mitochondria. DIO and Phalloidin/ A488 can be used to represent cellular areas in these studies. The trypsinized cells were seeded over DNA network-coated coverslips at a cell density of ∼20,000 cells/well and then incubated the cells for 24 hours at 37⁰C in 95% relative humid conditions with 5% CO_2_ levels.

#### Cellular adhesion and spreading

The RPE1 cells were grown on 10mm coverslips coated with 3WJ Y-type Hydrogel of varying polymeric concentrations. The coverslip, without any DNA polymer, acted as our control. The cells were split at 90% confluency and were seeded at 10,000 cells/well seeding density. The seeded cells were incubated for 24 hours, followed by fixing and permeabilizing the cells with 4% Paraformaldehyde (PFA) and 0.1% Triton X-100 solution. The cells were stained with Alexa-488 tagged phalloidin in 0.1% Triton X-100 solution at a 2.5 µg/mL working concentration. The coverslips were washed twice with 1x PBS (pH 7.4) and then mounted (inverted) over moviol with DAPI for fixation and counter-staining.

#### Mitochondrial Expression and Lipid profile change

The cells were cultured for 24 hours on DNA HG-coated coverslips for this experiment. After 24 hours of incubation, the old media was decanted, and the cells were washed with 1x PBS. The cells were stained with 2µg/mL of MitoTracker DeepRed (for staining mitochondria) and 2.5µg/mL of DIO (for staining Phospholipids) in DMEM serum-free media for 15 minutes. After this step, the coverslips were washed with 1xPBS, then fixed with 4% PFA, PBS washing, and mounting over moviol and DAPI. The cells were imaged at 405nm (DAPI), 488nm (DIO), and 633nm (MitoTracker DeepRed).

#### Endocytic Uptake Study of Cholera Toxin-B and Transferrin

After 24 hours of growth, the old media was decanted, and the cells were washed with 1x PBS. The cells were then incubated with 2µg/mL of CTxB and 5µg/mL of Tf in DMEM serum-free media for 15 minutes, 30 minutes, and 45 minutes, respectively, for multiple time points. The coverslips were washed with 1xPBS, followed by fixing the cells with 4% PFA, washing with PBS again, and then mounting the coverslip over moviol and DAPI. The cells were imaged at 405nm (DAPI), 488nm (CTxB), and 633nm (Tf).

#### DNA Hydrogels Influence Membrane Receptor Expression

The cells were first fixed with 4%PFA and treated with a blocking buffer for 1 Hour for the immunofluorescence studies. The cells were then incubated with Primary antibody (at 1:100 to 1:500 ratio in blocking buffer) for 2 Hours at 37°C followed by staining the primary Ab-tagged proteins of interest with Fluorophore-conjugated secondary antibody (∼ 1:1000 ratio in Blocking buffer) and kept for incubation at 37°C for 2 hours. The working dilutions of primary antibodies varied from proteins to proteins. The coverslips were then washed with 1x PBS before mounting them over moviol and DAPI mix. We used the same protocol for the CD98, CD147, Vinculin, β-integrin, and E-cadherin immunostaining. The images were taken on a Leica TCS SP5 confocal microscope with 63x objective (oil immersion type). DAPI was excited by a 405nm laser; for DIO and Phalloidin/Alexa 488 dye, a 488nm laser was used, and a 633nm laser was used to excite MitoTracker DeepRed dye. The images were processed and quantified on ImageJ software and the graph was plotted by GraphPad Prism software.

#### Cytotoxicity assay

The cytotoxicity of different DNA samples was tested by MTT (3-[4,5-dimethylthiazol- 2-yl]-2,5 diphenyl tetrazolium bromide) assay (Supplementary Figure 9-c). Five different DNA samples (T, Y, X, Pentamer, and hexamer) were taken in eight different concentrations (100 µM, 50 µM, 25 µM, 10 µM, 5 µM, 1 µM, 0.1 µM, and 0 µM) in triplicates were added into 96well plates and incubated overnight for this study. Then, the trypsinized cells were loaded in each well at a 10000 cells/well density and incubated for 24 hours at 37⁰C incubator with 5% CO_2_ and 95%RH levels. Remove the used media and wash the cells with PBS once. Then, add 100µL of MTT solution (0.5mg/mL) in DMEM-SF media into each well and incubate for 4 hours at 37⁰C. after 4 hours, remove the MTT solution gently without disturbing the formazan crystals. Add DMSO later to dissolve the crystal by gently shaking over a gel rocker for 5 minutes before taking its reading. The fluorescent signal of these formazan crystals can be acquired at 540nm and 720nm to remove the background noise. The fluorescent reading was taken on a Perkin Elmer multimode plate reader. The graph is plotted with GraphPad Prism software.

## Author Contributions

D.B. conceived the idea, and A.S. and D.B. planned the experiments. A.S. did the synthesis and characterization by EMSA, Bio-AFM, NTA, confocal microscopy, and cellular area change experiments. N.S. has contributed to this work with the endocytic uptake study and the confocal microscopic imaging. A.S. analyzed all the data, and it was cross-checked by all mentors. All the authors contributed to writing and reading the manuscript draft.

## Acknowledgments

The authors sincerely thank all of the members of the D.B. group for critically reading the manuscript and for their valuable feedback. The authors recognize IIT Gandhinagar’s infrastructure and financial support for the conduct of this research. NS and AS acknowledge the financial support from the Ministry of Education, Government of India. AS thanks PMRF fellowship. The authors acknowledge the Central Instrumentation Facility at IIT Gandhinagar for assistance with Field emission-SEM microscopy, AFM microscopy, and Confocal microscopy. D.B. thanks SERB, and GoI for the Core research grant, IITGN for the start-up grant, and Gujcost-DST, GSBTM, and STARS-MoES for research grants. D.B. is a member of the Indian National Young Academy of Sciences (INYAS).

