## Supporting information for "Programmable soft DNA hydrogels stimulate cellular endocytic pathways and proliferation"

### Supplementary Information

**Table 1:** Sequences of oligonucleotides used for the synthesis of DNA hydrogels

| Sr. No. | Name | Sequence |
|---|---|---|
| 1 | Y1 | 5'-CACGTGCGACCGATGAATAGCGGTCAGATCCGTACCTACTCG-3’ |
| 2 | Y2 | 5'-CACGTGCGAGTCGTTCGCAATACGACCGCTATTCATCGGTCG-3’ |
| 3 | Y3 | 5'-CACGTGCGAGTAGGTACGGATCTGCGTATTGCGAACGACTCG-3’ |
| 4 | T1 | 5'-CACGTGCGACAGCTGACTAGAGTCACGACCTGTACCTACTCG-3' |
| 5 | T2 | 5'-CACGTGCGAGTCGTTCTCAAGACGTAGCTAGGACTCTAGTCAGCTGTCG-3' |
| 6 | T3 | 5'-CACGTGCGAGTAGGTACAGGTCGTCGTCTTGAGAACGACTCG-3’ |
| 7 | X1 | 5'-CACGTGCGACCGATGAATAGCGGTCAGATCCGTACCTACTCG-3' |
| 8 | X2 | 5'-CACGTGCGAGTAGGTACGGATCTGCGTATTGCGAACGACTCG-3' |
| 9 | X3 | 5’-CACGTGCGAGTCGTTCGCAATACGGCTGTACGTATGGTCTCG-3' |
| 10 | X4 | 5'-CACGTGCGAGACCATACGTACAGCACCGCTATTCATCGGTCG-3' |
| 11 | 5WJ A1 | 5’-TGTCACTCGCAGAGCTTACCGGTGACGTTCCGC-3’ |
| 12 | 5WJ A2 | 5’-TGTCACTCGCAGAGCTTCCGCGTCCTCACCGGT-3’ |
| 13 | 5WJ A3 | 5’-TGTCACTCGCAGAGCTTGCCACGCTTTGGACGCGG-3’ |
| 14 | 5WJ A4 | 5’-TGTCACTCGCAGAGCTTGCGAGTGCAAAGCGTGGC-3’ |
| 15 | 5WJ A5 | 5’-TGTCACTCGCAGAGCTTGCGGAACGAAGCACTCGC-3’ |
| 16 | 5 WJ B1 | 5’-GCTCTGCGAGTGACATTACCGGTGACGTTCCGC-3’ |
| 17 | 5 WJ B2 | 5’-GCTCTGCGAGTGACATTCCGCGTCCTCACCGGT-3’ |
| 18 | 5 WJ B3 | 5’-GCTCGTCGAGTGACATTGCCACGCTTTGGACGCCG-3’ |
| 19 | 5 WJ B4 | 5’-GCTCTGCGAGTGACATTGCGAGTGCAAAGCTTGGC-3’ |
| 20 | 5 WJ B5 | 5’-GCTCTGCGAGTGACATTGCGGAACGAAGCACTCGC-3’ |
| 21 | 6 WJ A1 | 5’-TGTCACTCGCAGAGCTTCCATCTCACGTCCGCTAACCTAACGCCGACTTGG-3’ |
| 22 | 6 WJ A2 | 5’-TGTCACTCGCAGAGCTTCGTTCTGAGCACGAGTAGCGGACGTGAGATGG-3’ |
| 23 | 6 WJ A3 | 5’-TGTCACTCGCAGAGCTTCGCACCACCGTAAGCCAAACTCGTGCTCAGAACG-3’ |
| 24 | 6 WJ A4 | 5’-TGTCACTCGCAGAGCTTCCTAGTCTCGACCTGCGGCTTACGGTGGTGCG-3’ |
| 25 | 6 WJ A5 | 5’-TGTCACTCGCAGAGCTTCCTACACACGGTATCAAAGCAGGTCGAGACTAGG-3’ |

| | | |
|---|---|---|
| 26 | 6 WJ A6 | 5’-TGTCACTCGCAGAGCTTCCAAGTCGGCGTTAGGTGATACCGTGTGTAGG-3’ |
| 27 | 6 WJ B1 | 5’-GCTCTGCGAGTGACATTCCATCTCACGTCCGCTCCTAACGCCGACTTGG-3’ |
| 28 | 6 WJ B2 | 5’-GCTCTGCGAGTGACATTCGTTCTGAGCACGAGTTTAGCGGACGTGAGATGG-3’ |
| 29 | 6 WJ B3 | 5’-GCTCTGCGAGTGACATTCGCACCACCGTAAGCCACTCGTGCTCAGAACG-3’ |
| 30 | 6 WJ B4 | 5’-GCTCTGCGAGTGACATTCCTAGTCTCGACCTGCTTGGCTTACGGTGGTGCG-3’ |
| 31 | 6 WJ B5 | 5’-GCTCTGCGAGTGACATTCCTACACACGGTATCAGCAGGTCGAGACTAGG-3’ |
| 32 | 6 WJ B6 | 5’-GCTCTGCGAGTGACATTCCAAGTCGGCGTTAGGTTTGATACCGTGTGTAGG-3’ |

**Supplementary figures:**

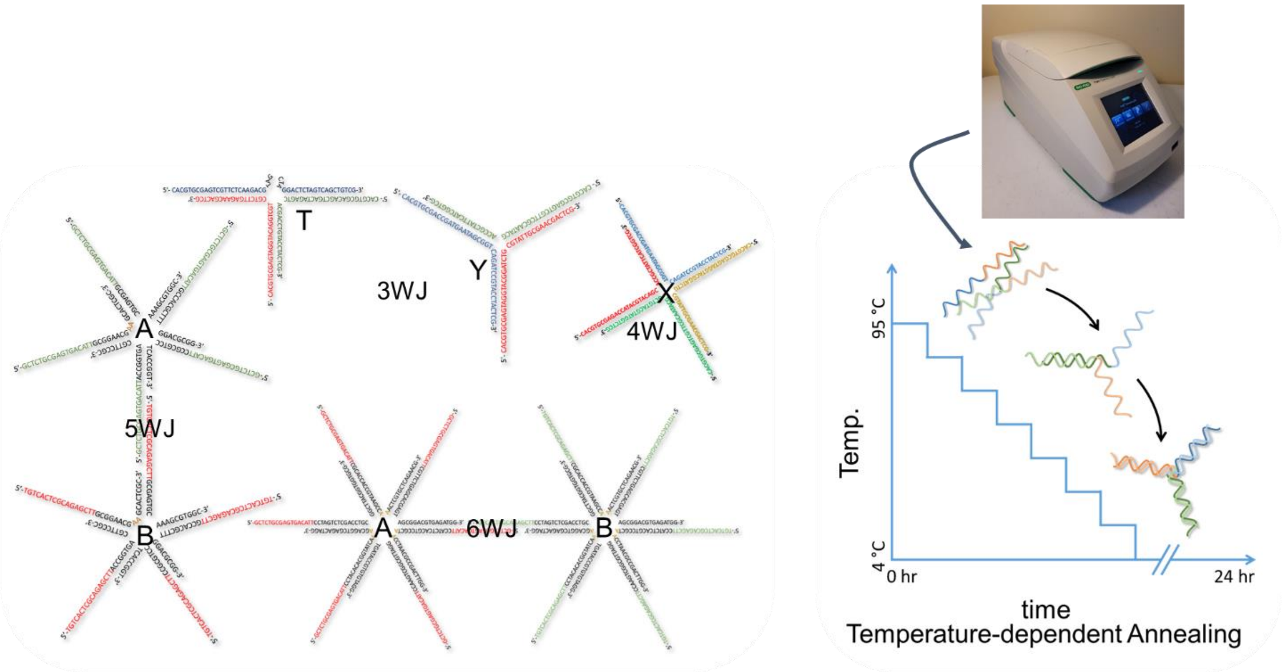


***Supplementary Figure 1:*** *Schematics of self-assembled DNA Supramolecular branched structures. The left side shows the temperature-dependent self-annealing property of ssDNA to form higher-order complexes.*

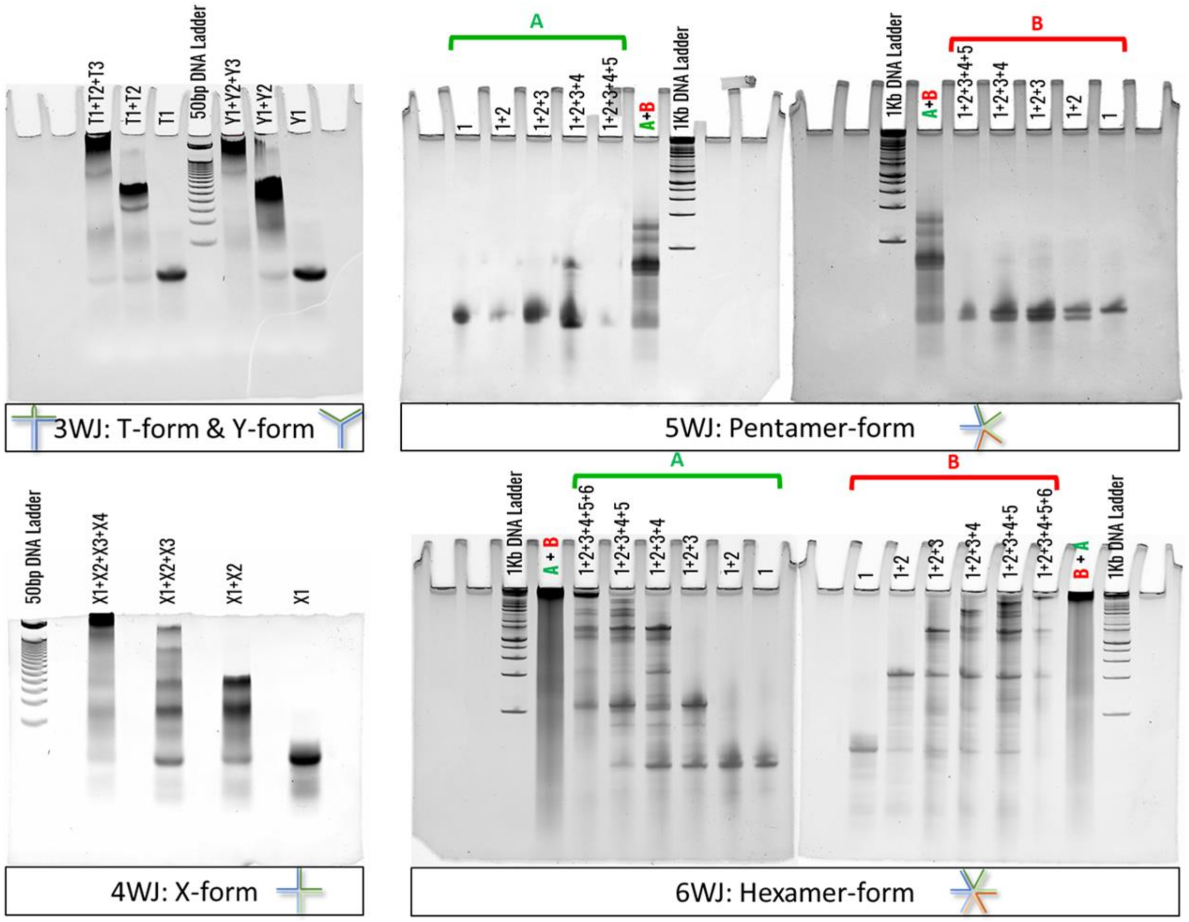


**Supplementary Figure 2:** Electrophoretic Mobility Shift Assay (EMSA) of subsequent strands of DNA in forming higher order complex structures. The ladderlike band patterns suggest an increase in the molecular size of the assembled structures.

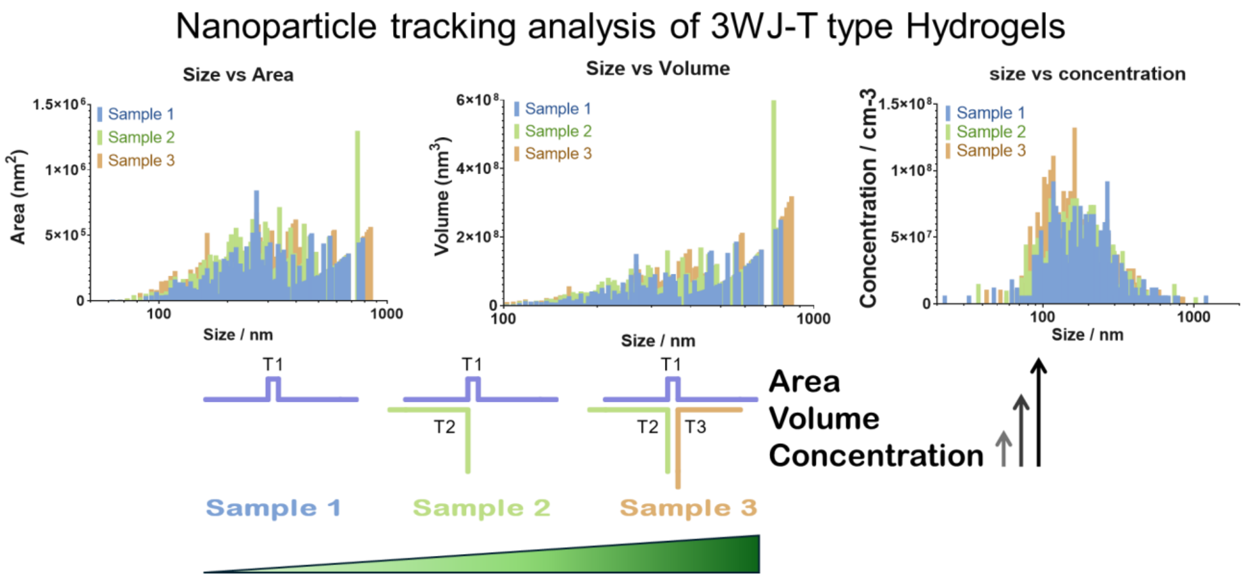


***Supplementary Figure 3:*** *NTA-based formation of higher order complexes in subsequent samples of 3WJ T-type DNA strands. The graph shows an increment in the Area, Volume, and concentration of higher-order complexes forming by self-assembly of all three strands of 3WJ T-type DNA Hydrogels, namely T1+T2+T3.*

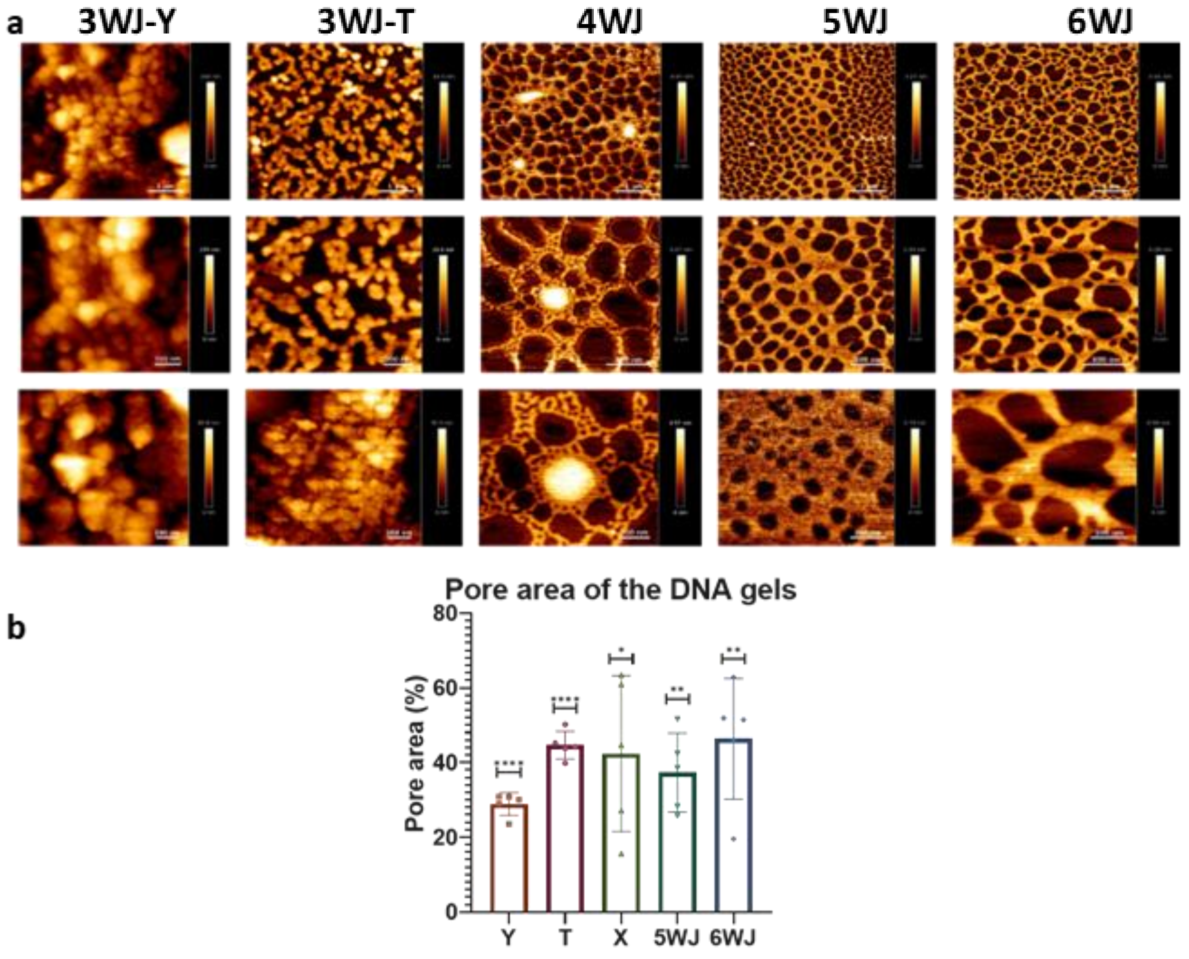

**Supplementary Figure 4**: Morphological characterization of DNA Hydrogels at 100uM DNA concentrations using AFM (a) Images were acquired at 5*5 μm2 (Top), 2.5*2.5 μm$^2$ (Middle), and 1*1 μm$^2$ (Bottom), with scale bars of 1μm, 500nm, and 200nm, respectively. (b) The porosity of the gel was back calculated from the void spaces present within the AFM by applying a threshold and measuring the area of the pores. N=5 images, ***, signifies p-value= 0.0003 (Ordinary One-way ANOVA).

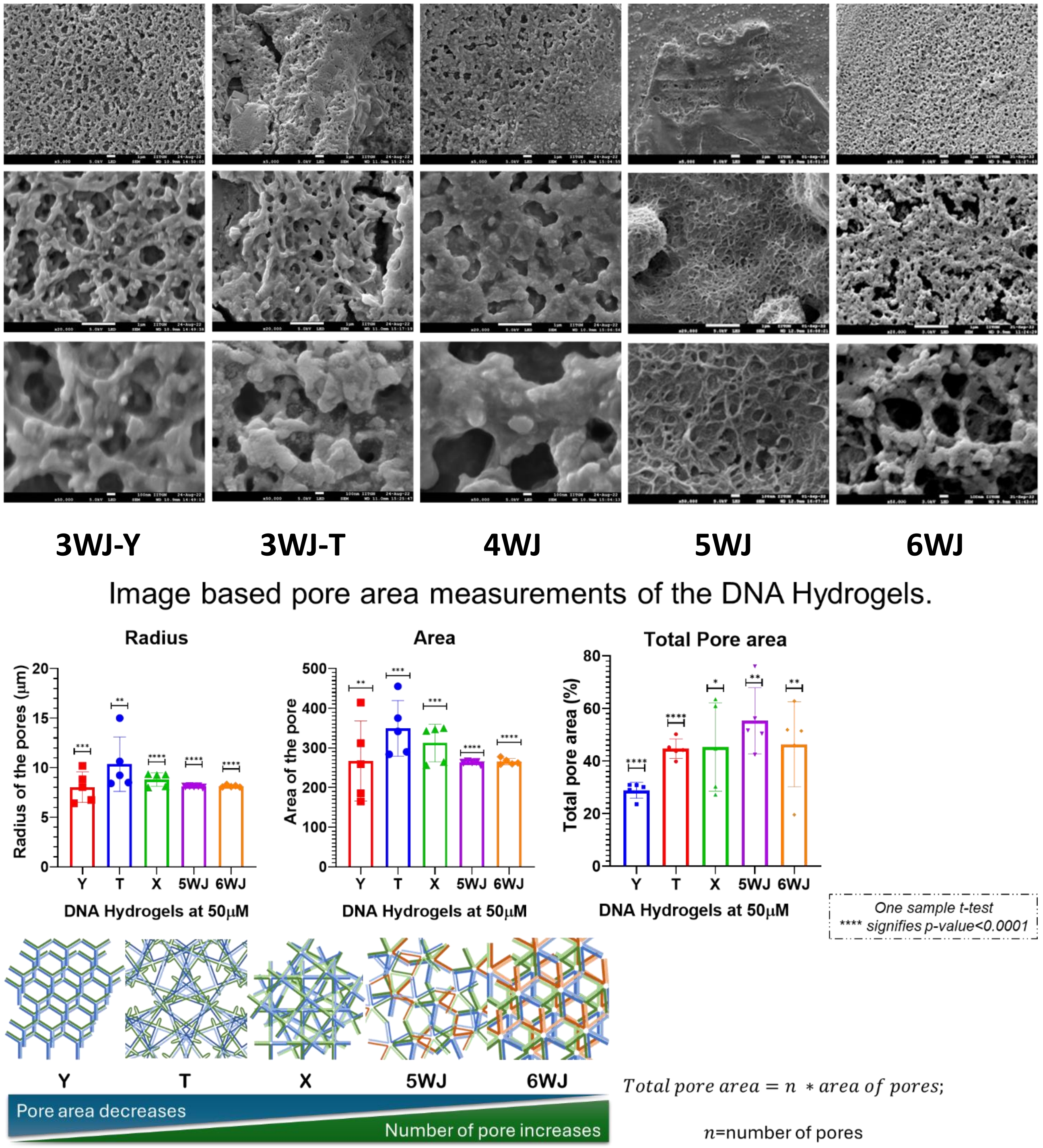


**Supplementary Figure 5**: Morphological characterization of DNA Hydrogels at 100uM DNA concentrations using FE-SEM (a) Images were acquired at 5000x (Top), 20000x (Middle), and 50000x (Bottom) magnifications, with scale bars of 1μm, 1 μm, and

100nm, respectively. (b) The porosity of the gel was back-calculated from the void spaces present within the SEM images by applying a threshold and measuring the area of the pores. N=5 images, ***, signifies p-value= 0.0003 (Ordinary One-way ANOVA).

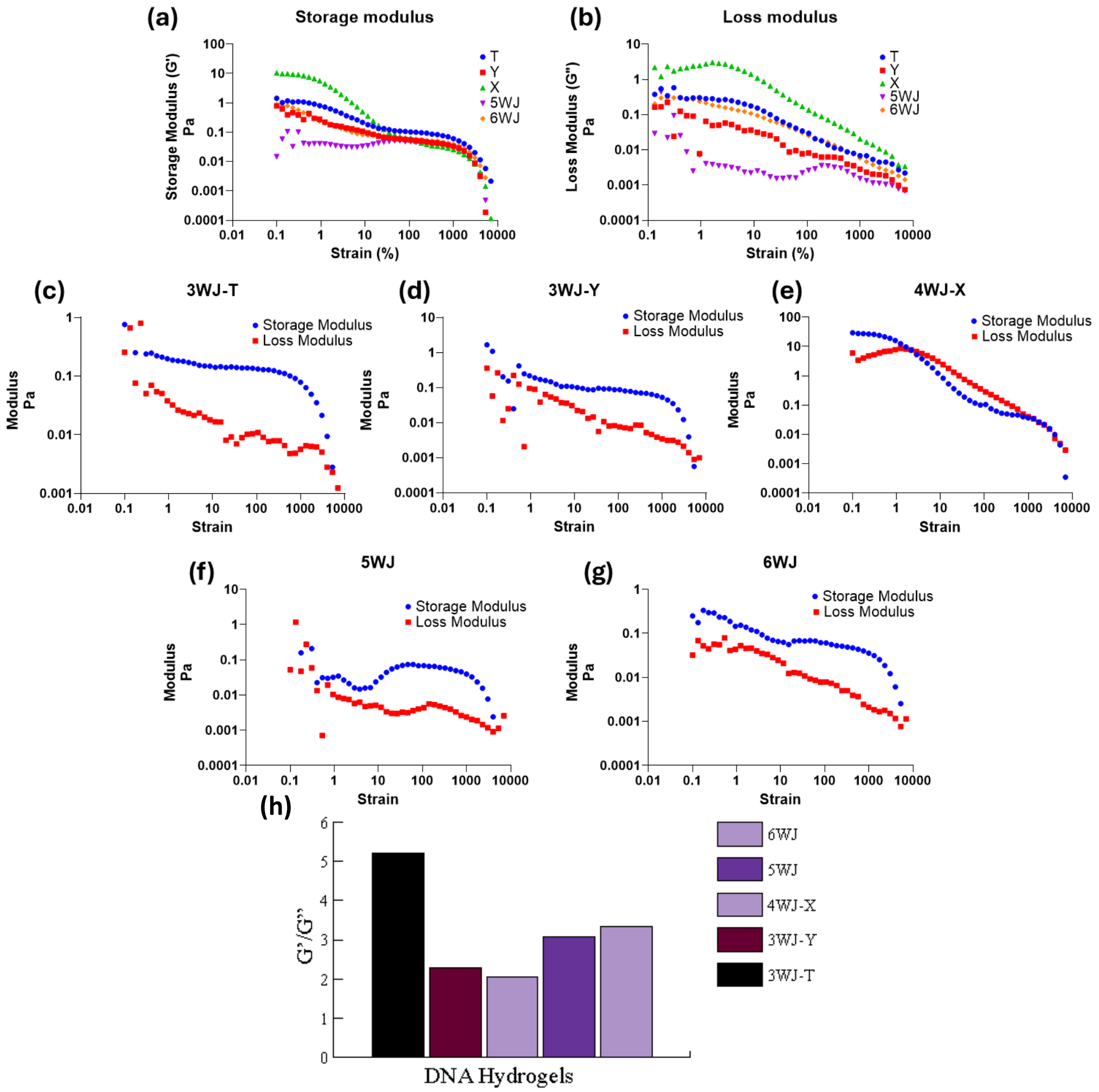


**Supplementary Figure 6:** Rheology of the DNA hydrogels at 50uM. The Oscillatory Strain sweep test of these hydrogels shows that these polymeric networks behave as gels as their storage modulus is higher than loss modulus values. (a) Storage modulus and (b) loss modulus of all the DNA Hydrogel candidates at 50uM. The exact modulus values of 3WJ-T (c), 3WJ- Y (d), 4WJ-X (e), 5WJ-Pentamer (f), and 6WJ-Hexamer (g), have been shown as distinct graphs, respectively. The strain sweep test was performed at 41 measurement points, keeping angular frequency, ω= 6.28 rad/sec, and Temp, T=25°C constant. (h) the ratio of storage (G') modulus and the loss (G") modulus was calculated for all DNA HG samples at 1% oscillatory strain and 1Hz frequency.

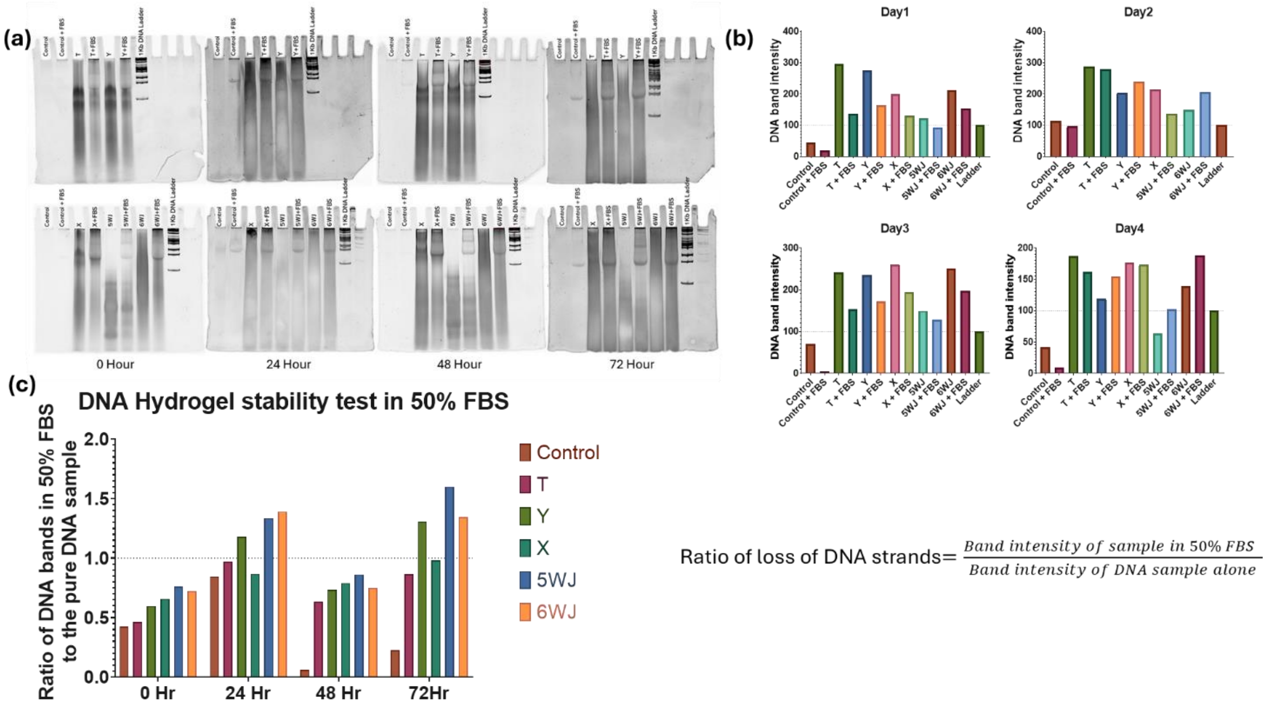


***Supplementary Figure 7:*** *DNA Hydrogel Stability in 50% FBS was tested by incubating the DNA HG samples for up to 72 hours, and the integrity of the DNA bands was checked on 10% Native PAGE. (a) Gel electrophoretic bands of DNA hydrogel samples and DNA Hydrogel samples mixed in 50% FBS. The gel was 10% Native PAGE. (b) quantified measurements of the DNA bands for each condition normalized against the intensity of the 1Kb DNA ladder for a period of four days. (c) The ratio of band intensities of DNA samples in FBS to the DNA sample alone shows the degree of degradation over a period of 72 Hours. A ratio higher than one indicates degradation of DNA strands from pure DNA samples, while ratios lower than one denotes degradation of DNA strands in the FBS sample. Each lane's band intensity was calculated by plotting a graph in ImageJ software, and calculating the area under the curve, providing the total intensity of the entire lane. Comparing the lane intensities with the FBS sample counterparts shows degree of DNA strand degradation.*

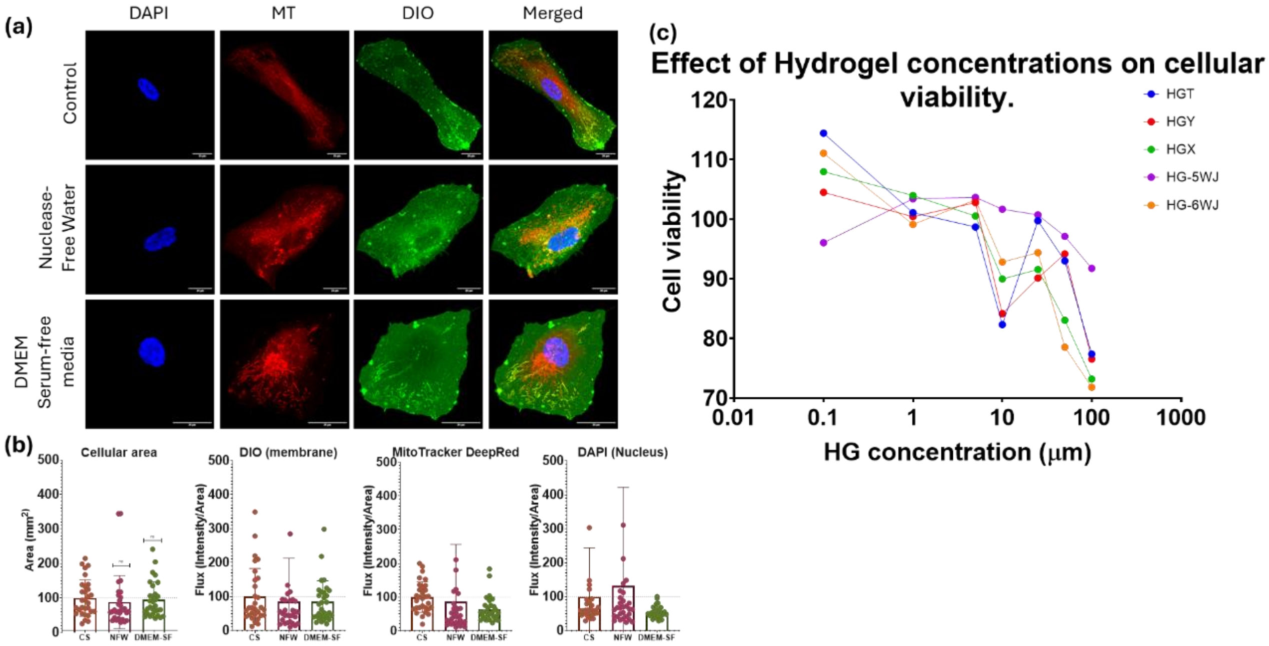


**Supplementary Figure 8:** Stability of DNA hydrogels within cation rich DMEM media. (a) Cells were grown over 3WJ-Y type DNA Hydrogels at 10uM concentration for 24 Hours, followed by staining with DAPI (Blue), MT (Red), and DIO (Green). Divalent cation-rich cell-culture grade DMEM media was used to test the stability of the DNA hydrogels. (b) No significant difference was observed by changing the solvent on the Area of cells, DIO, DAPI, and MT expression patterns. n=30; ns, signifies p-value <0.05. (c) MTT assay of RPE1 cells grown over Different DNA hydrogels shows increased cellular viability with decreased hydrogel concentration. The readings of 50uM Hydrogel show the same pattern from the previous confocal analysis, thereby increasing the confidence of this data obtained. At 50uM, the maximum cellular viability can be seen in 5WJ, and the minimum was observed in 6WJ. N=3

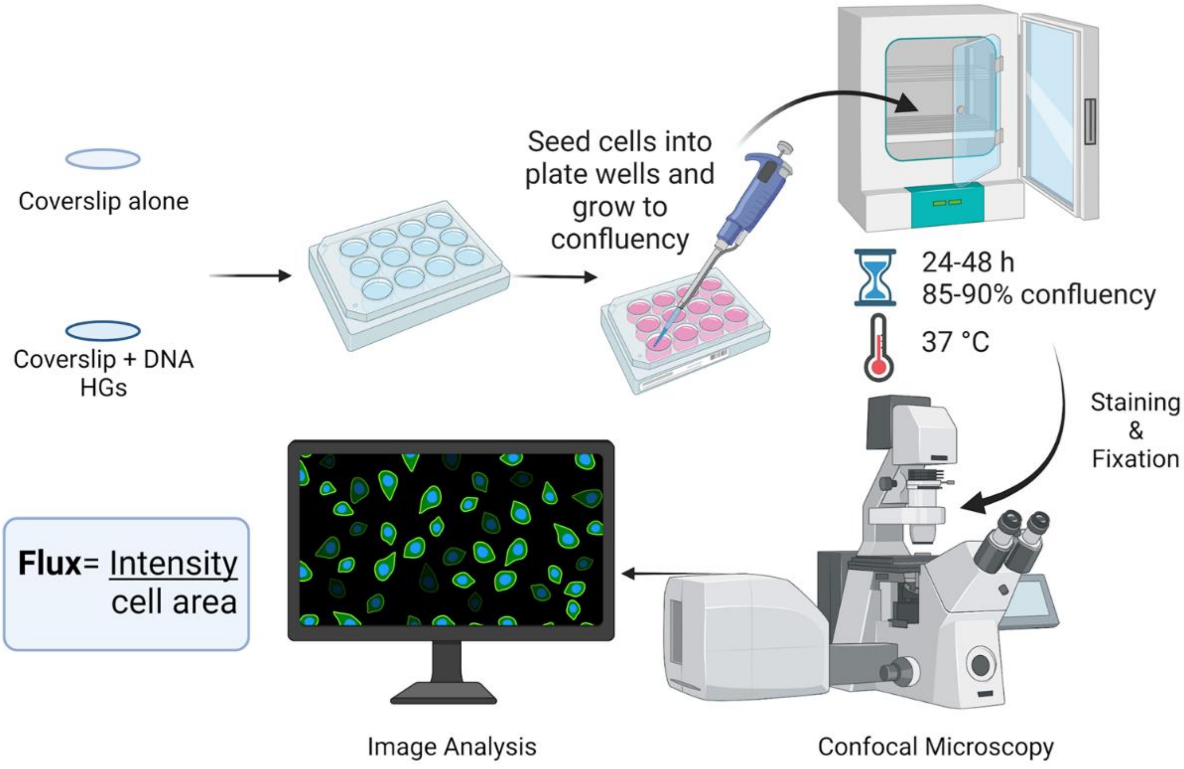

**Supplementary Figure 9:** Schematics of the experimental plan for cell culture applications. The coverslips were coated with DNA hydrogels, followed by overnight incubation at RT. The cells were then seeded onto the coverslips (both coated and uncoated) within the cell-culture well plates. The cells were then allowed to grow for 24 hours, followed by fluoro-labeling/ immunolabelling of the target of interest and fixing the cells. The fixed cells were then imaged under a confocal microscope, followed by image analysis to observe the expression pattern.

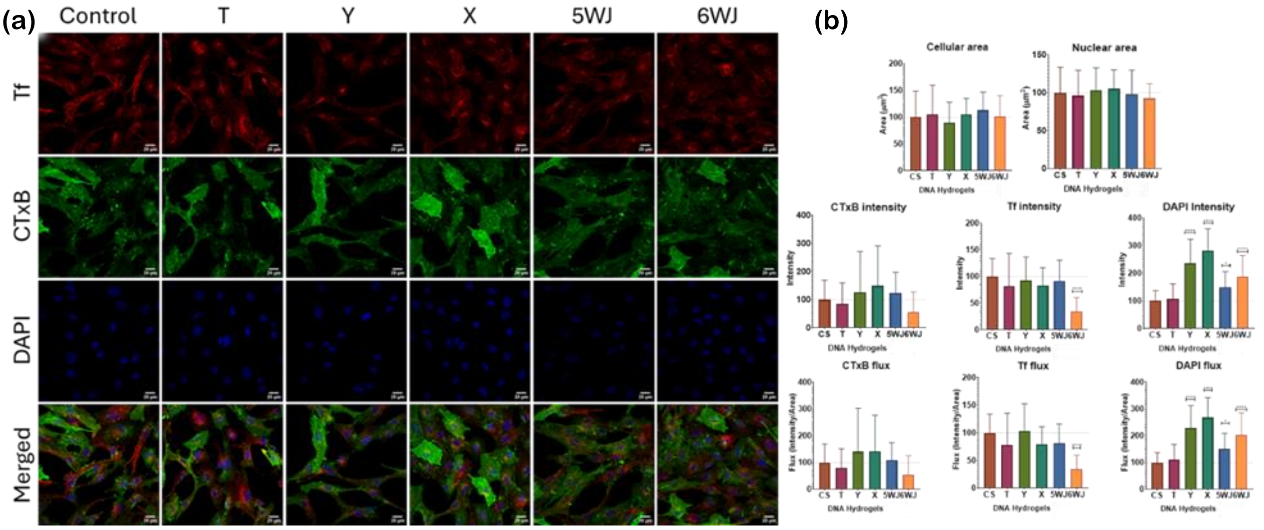


**Supplementary Figure 10:** (a,b): Endocytic Uptake study of Cholera Toxin-B (CTxB-FITC; Green), Transferrin (Tf-Alexa 647; Red) and DAPI (Blue). (a) Confocal images showing the relative expression pattern of CTxB, Tf, and DAPI within RPE-1 cells when grown over different Hydrogels of 50µM Concentrations, followed by an incubation period of 30 minutes. Scale bar =20 µm. (b) Graphs showing quantified cellular area, nuclear area, Raw Intensity, and Flux of CTxB, TF, and DAPI, respectively. Statistics: **** signifies p-value < 0.0001, n=30 (One-way ordinary ANOVA). (c-d) show endocytic Uptake in RPE1 cells after 30 mins of incubation post-treatment.

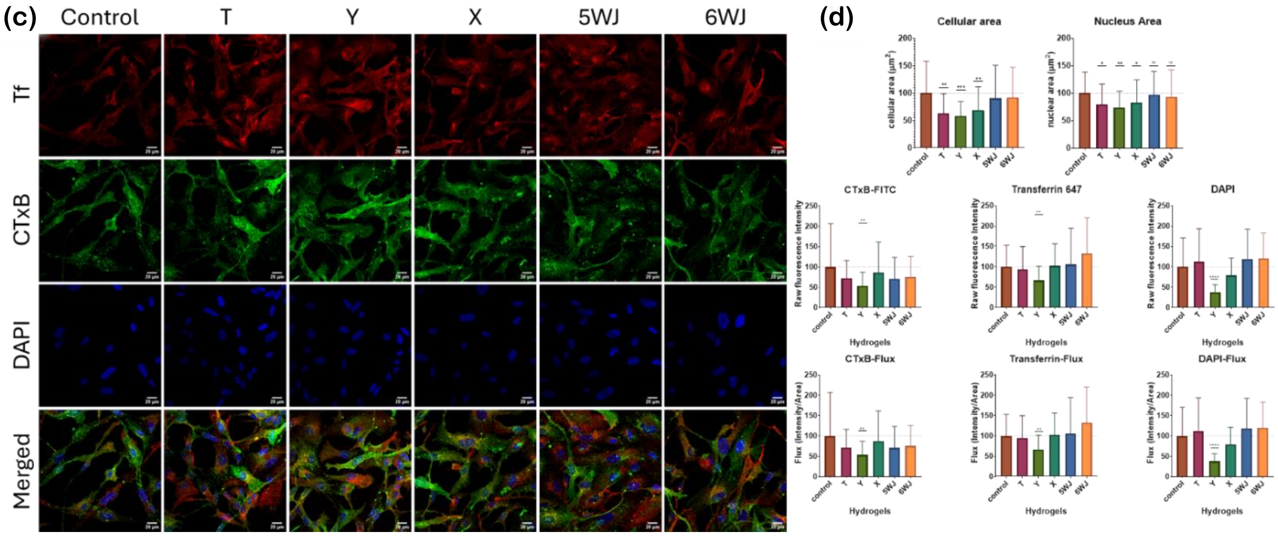

**Supplementary Figure 10:** (c,d): Endocytic Uptake study of Cholera Toxin-B (CTxB-FITC; Green), Transferrin (Tf-Alexa 647; Red) and DAPI (Blue). (c) Confocal images showing the relative expression pattern of CTxB, Tf, and DAPI within RPE-1 cells when grown over different Hydrogels of 50μM Concentrations, followed by an incubation period of 45 minutes. Scale bar =20 μm. (d) Graphs showing quantified cellular area, nuclear area, Raw Intensity, and Flux of CTxB, TF, and DAPI, respectively. Statistics: **** signifies p-value < 0.0001, n=30 (One-way ordinary ANOVA). (c-d) show endocytic Uptake in RPE1 cells after 45 mins of incubation post-treatment.

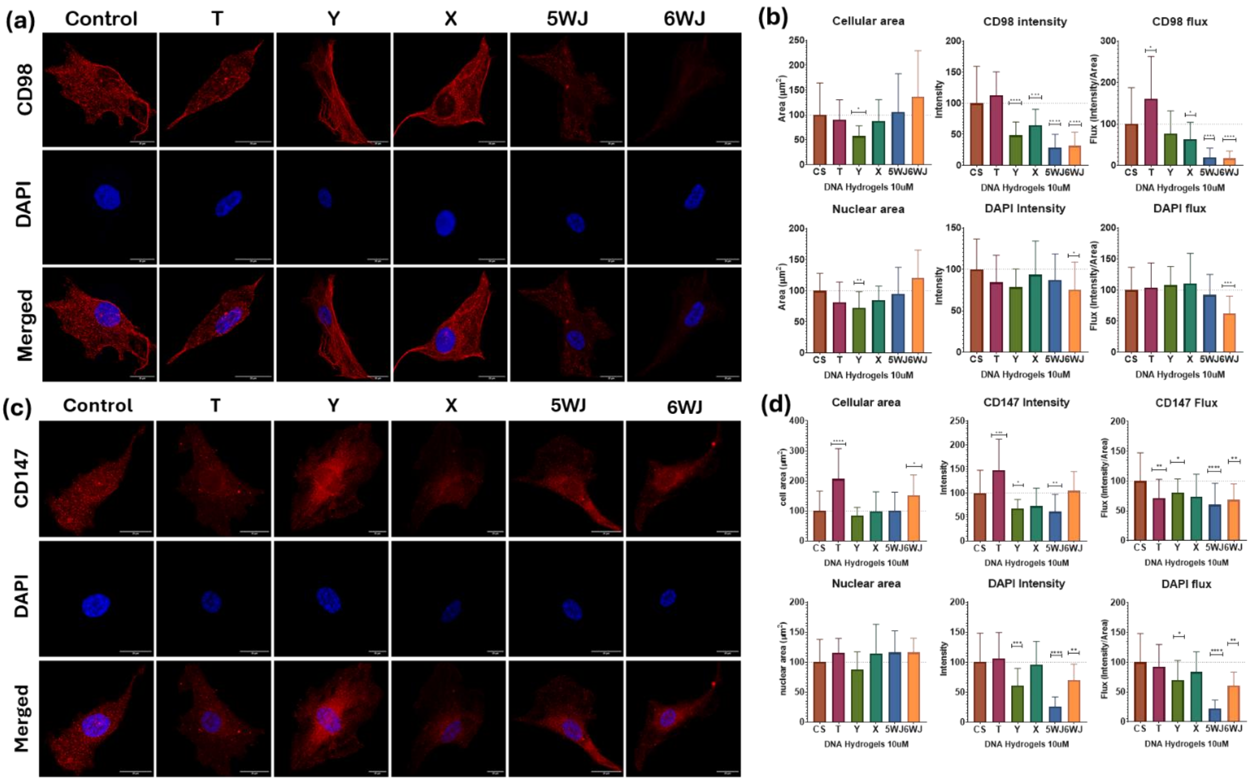


**Supplementary Figure 11:** Immunofluorescence study of CD98 (a-b) and CD147 (c-d) expression levels in RPE 1 cells grown over 10 μM of DNA Hydrogels. (a,b) Confocal microscopic images of RPE 1 cells stained with CD98 (Red) and DAPI (Blue). (c,d) Quantified data shows changes in the cellular area, nucleus area, intensity, and flux of CD147 and DAPI, respectively. N = 30 cells; **** signifies p-value< 0.0001 (Ordinary One-way ANOVA). The scale bar is 20um.

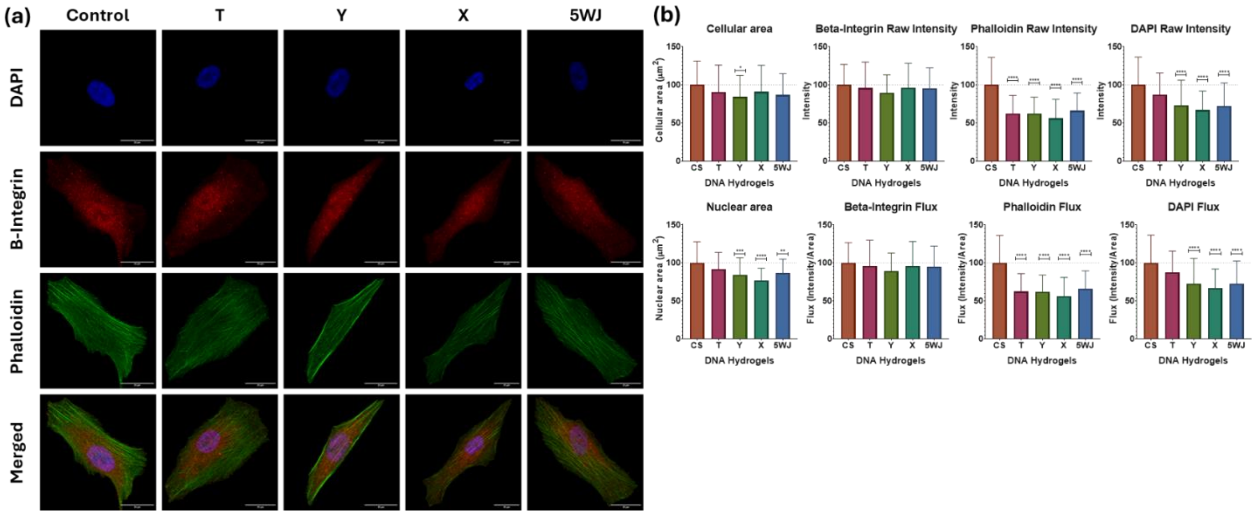


***Supplementary Figure 12:*** *Immunofluorescence staining of β –Integrin, phalloidin, and DAPI in RPE 1 cells grown over 50uM of DNA Hydrogels. (a) Confocal microscopic images of RPE 1 cells stained with β –Integrin (Red), Phalloidin (Green), and DAPI (Blue). The scale bar is 20um. (b) Quantified data showing changes in the cellular area, nucleus area, Intensity, and flux changes of β –Integrin, Phalloidin, and DAPI, respectively. N = 30 cells; ****, signifies p-value < 0.0001 (Ordinary One-way ANOVA).*

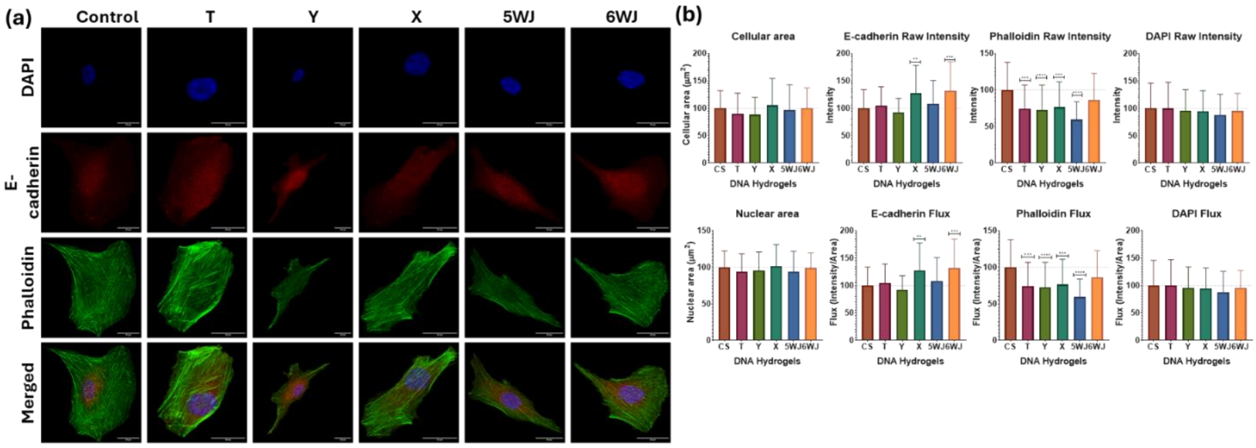


**Supplementary Figure 13:** Immunofluorescence staining of E-cadherin, phalloidin, and DAPI in RPE1 cells grown over 50uM of DNA Hydrogels. (a) Confocal microscopic images of RPE 1 cells stained with E-cadherin (Red), Phalloidin (Green), and DAPI (Blue). The scale bar is 20um. (b) Quantified data shows changes in the cellular area, nucleus area, intensity, and flux changes of E-cadherin, Phalloidin, and DAPI, respectively. N = 30 cells; ****, signifies p-value< 0.0001 (Ordinary One-way ANOVA).
